# The Biosynthetic Pathway to the Pyrroloiminoquinone Marine Natural Product Ammosamide C

**DOI:** 10.1101/2025.09.03.674030

**Authors:** Josseline S. Ramos-Figueroa, Lingyang Zhu, Matthew Halliman, Wilfred A. van der Donk

**Affiliations:** Department of Chemistry and Howard Hughes Medical Institute, University of Illinois at Urbana-Champaign, Urbana, Illinois 61801, USA; School of Chemical Sciences NMR Laboratory, University of Illinois at Urbana-Champaign, Urbana, 61801, IL, USA

## Abstract

Ammosamide C is a marine natural product containing a highly decorated pyrroloiminoquinone core. Studies on the biosynthetic gene cluster (BGC) that produces ammosamides previously revealed that they are made by a series of posttranslational modifications (PTMs). The BGC includes genes encoding a precursor peptide AmmA and four enzymes known as PEptide Aminoacyl-tRNA Ligases (PEARLs). Initial studies into the ammosamide biosynthetic pathway demonstrated Trp addition to a precursor peptide by the PEARL AmmB_2_. Thereafter, sequential modifications by several enzymes including two other PEARLs lead to the formation of a peptide intermediate bearing a C-terminal diaminoquinone. In the present work, we present the biosynthetic steps that convert this intermediate to ammosamide C. The PEARL AmmB_4_ unexpectedly appends an arginine to the C-terminus of the aforementioned intermediate. Then, C-terminal proteolysis by the heterodimeric TldD/E-like protease Amm12/13 releases a dipeptide, which is subsequently cleaved by the dipeptidase Amm19 to produce a Trp-derived diaminoquinone. Amm3 next catalyzes the conversion of this Trp derivative to the corresponding chlorinated ammosamaic acid. Finally, Amm23 methylates this intermediate and a putative aminotransferase Amm20 performs an amidation to arrive at ammosamide C; the order of these last two steps could not be determined. This study reveals an unexpectedly lengthy route to ammosamide that illustrates the opportunistic nature of natural product biosynthesis, demonstrates a role for a PEARL that is unlike previous roles, identifies steps that are not PTMs, and adds Arg-tRNA to the growing repertoire of amino acyl tRNAs that are used by PEARLs.

## Introduction

Marine natural products represent a largely untapped pool of chemically diverse, bioactive molecules.^1,2^ Pyrroloiminoquinones are a subgroup of marine natural products known for their potent and wide ranging bioactivities such as antifungal, antitumor, antiviral, and antimicrobial activities.^3,4^ Structurally, pyrroloiminoquinone-containing alkaloids are composed of a pyrrolo[4,3,2-*de*]quinoline core, which is believed to impart antiproliferative and cytotoxic properties against several cancer cell lines. Pyrroloiminoquinone derived natural products isolated thus far include the makaluvamines, discorhabdins, and ammosamides.^3,4^ Because of their proven biological effects, development of concise synthetic routes to pyrroloiminoquinones have been achieved,^5,6^ but understanding of their biosynthesis is still in an early stage.

In 2009, Fenical and coworkers isolated the pyrroloiminoquinone-derived ammosamides A and B from *Streptomyces* sp. CNR-698 exhibiting anticancer activities against several cancer cell lines and targeting myosin.^7,8^ With the advancement of whole genome sequencing and bioinformatic approaches, the BGC that produces the ammosamides was identified in 2016.^9,10^ Through heterologous expression of the *amm* BGC (Figure 1B), an additional pyrroloiminoquinone-containing metabolite accumulated early in the fermentation termed ammosamide C (Figure 1A),^10^ which is likely the true natural product and may have an electrophilic mechanism of action.^11^ Bioinformatic analysis of the BGC revealed a set of genes encoding putative truncated lantibiotic dehydratases, proteases, as well as a short peptide sequence, resembling a BGC for a ribosomally synthesized and posttranslationally-modified peptide (RiPP).^12^ In addition, the ammosamide BGC displayed homology in gene sequence and architecture to the BGC encoding the mTOR inhibitor lymphostin that also contains a pyrroloiminoquinone core (Figure 1A).^13,14^ Through genetic manipulations, the formation of ammosamide A-C was shown to be strictly dependent on the encoded short peptide AmmA, the truncated lantibiotic dehydratases AmmB_1_, AmmB_2_, AmmB_3_, and AmmB_4_, as well as the peptidases Amm19, Amm12 and Amm13 (Figure 1B).^10^ Moreover, deletion of the genes encoding several other enzymes such as the methyltransferase Amm23 and chlorinase Amm3 were shown to accumulate late intermediates in the pathway including desmethyl ammosamide and ammosamaic acid (Figure S1).

**Figure 1.**
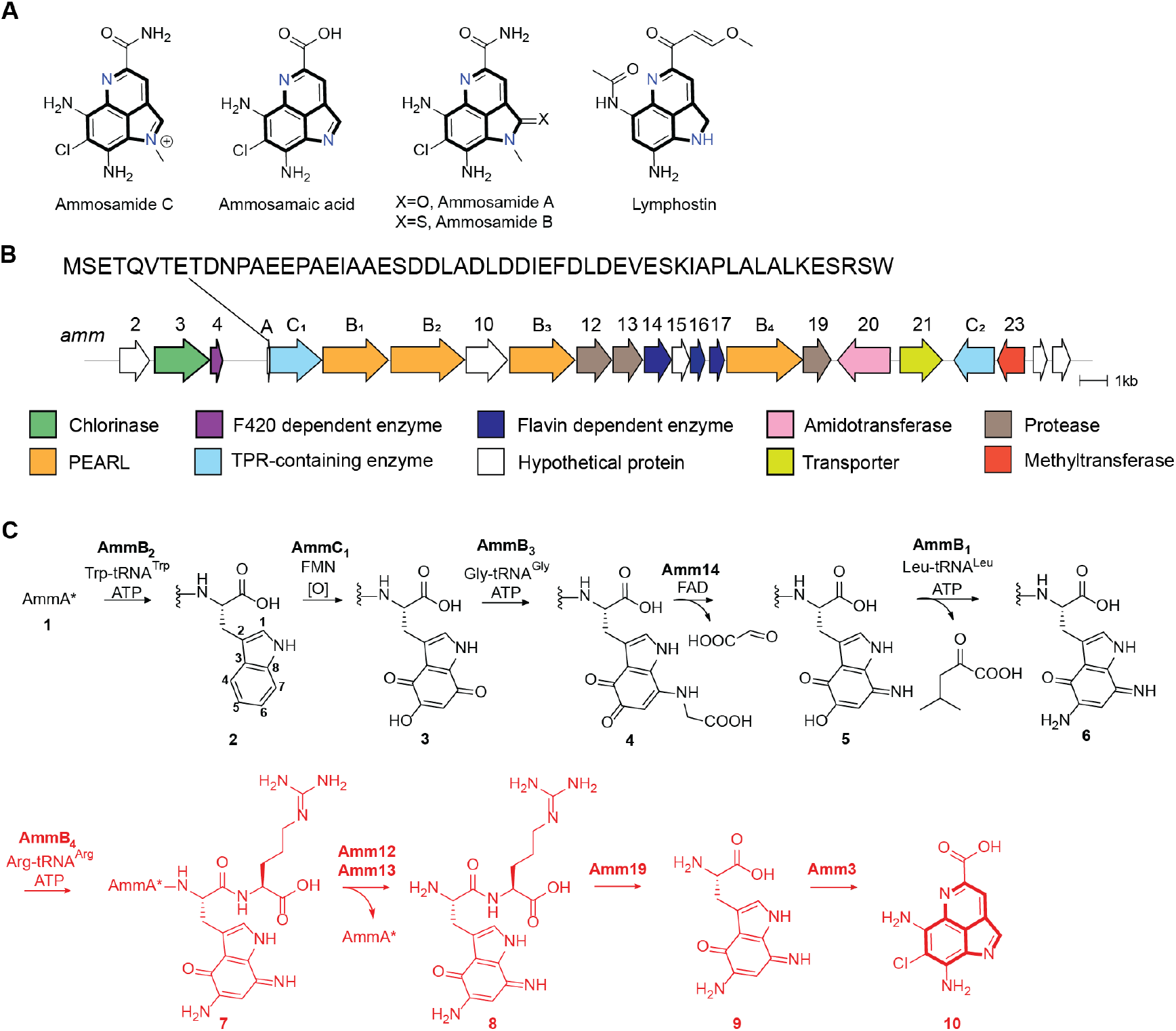
(A) Pyrroloimiquinone-containing marine natural products with the common scaffold highlighted in bold. (B) Ammosamide biosynthetic gene cluster identified in the genome of *Streptomyces* sp. CNR698.^10^ The sequence of the precursor peptide AmmA is shown; Two steps convert AmmA into AmmA* (for the sequence of AmmA*, see Figure S1).^15^ (C) Previously characterized steps in the biosynthetic pathway towards the formation of Ammosamide C. In red are the steps characterized in this work.

Independently, truncated lantibiotic dehydratases (Pfam PF04738) were shown to catalyze the appendage of amino acids to the C-termini of scaffold peptides in an amino acyl-tRNA and ATP dependent fashion and were termed PEptide Aminoacyl-tRNA Ligases (PEARLs).^15^ The *amm* BGC encodes four of these PEARLs (Figure 1B) and initial work showed that the PEARL AmmB_2_ catalyzes the addition of Trp to the C-terminus of a ribosomally produced peptide (Figure 1C). Thereafter, trihydroxylation by AmmC_1_ followed by oxidation (possibly by the quinone reductase Amm17)^16^ produced the corresponding quinone intermediate **3**, which was processed by AmmB_3_ by appending a glycine to the C7 position of the indole core (Figure 1C, **4**).^17^ The oxidoreductase Amm14 then catalyzed the oxidation of the glycyl-indole imine adduct producing the corresponding aminoquinone **5** and glyoxylate.^16^ Another amino transfer event to position C5 of the indole was catalyzed by the PEARL AmmB_1_ and used Leu-tRNA rather than Gly-tRNA as nitrogen donor to produce diaminoquinone **6**.^16^ The previous work left fourteen genes in the BGC uncharacterized that are presumably involved in transforming **6** to ammosamide C.

In this work, the subsequent steps towards the formation of ammosamide C were elucidated and the enzymes characterized. First, the final PEARL in the BGC, AmmB_4_, catalyzes arginine appendage to intermediate **6** to produce **7** (Figure 1C). Thereafter, proteolytic cleavage is catalyzed by the TldD/E-like heterodimeric Amm12/Amm13 protease releasing dipeptide **8**. The dipeptidase Amm19 then liberates the aminoquinone modified Trp (**9**). Chlorination by the flavin-dependent halogenase Amm3 results in the formation of the chlorinated ammosamaic acid, **10**. Finally, the methyltransferase Amm23 is shown to catalyze the methylation of this intermediate. This work expands the examples of pathways in which nature uses a scaffold peptide to build complexity on a peptide backbone and then through proteolysis cleaves off the decorated bioactive molecule to restart the biosynthesis on the scaffold peptide. In addition, this study illustrates new aspects of these pathways such as further tailoring after the posttranslationally generated structure is removed by proteolysis, and incorporation of an amino acid (Arg) that does not end up in the final product and appears to function for recognition by subsequent enzymes to prevent premature termination of the pathway. With the addition of Arg-tRNA as a PEARL substrate, the total number of aminoacyl tRNAs that have been shown to be used by these enzymes comes to eight, and AmmB_4_ is the first example that adds a charged amino acid. This expansion will benefit efforts that may allow future prediction of the amino acid specificity of uncharacterized PEARLs.

## Results and discussion

### AmmB_4_ catalyzes the appendage of Arg to the C-terminus of diaminoquinone intermediate 6

To continue the elucidation of the complete *amm* pathway, we again used the co-expression methodology that had been successful in identifying each consecutive step up to intermediate **6**.^16,17^ Coexpression of AmmB_4_ using an *E. coli* codon-optimized gene with the His-tagged precursor peptide AmmA* (Figure S1) in addition to all modifying enzymes required to form intermediate **6** resulted in a new product as demonstrated by matrix-assisted laser desorption/ionization time-of-flight (MALDI-TOF) mass spectrometry (MS) analysis (Figure 2). The ion for the diaminoquinone intermediate **6** disappeared and a new peak with a mass shift of +156 Da was observed. High-resolution MS/MS analysis after trypsin digestion was consistent with the increase in mass corresponding to the addition of an Arg residue to the C-terminus of the peptide chain (Figure 2C). To confirm this assignment, we purified the peptide from large scale expression (>80 L of culture) by high performance liquid chromatography (HPLC), digested the peptide with chymotrypsin, and performed 2D NMR experiments on the resulting C-terminal tripeptide. As expected, the Arg addition was confirmed to have occurred at the C-terminal carboxylate as shown by TOCSY, NOESY and HSQC experiments. Briefly, correlations between the NH proton (δ 7.58 ppm) of the Arg to the α proton (δ 4.50 ppm) of the modified Trp were observed (Figure S2).

**Figure 2.**
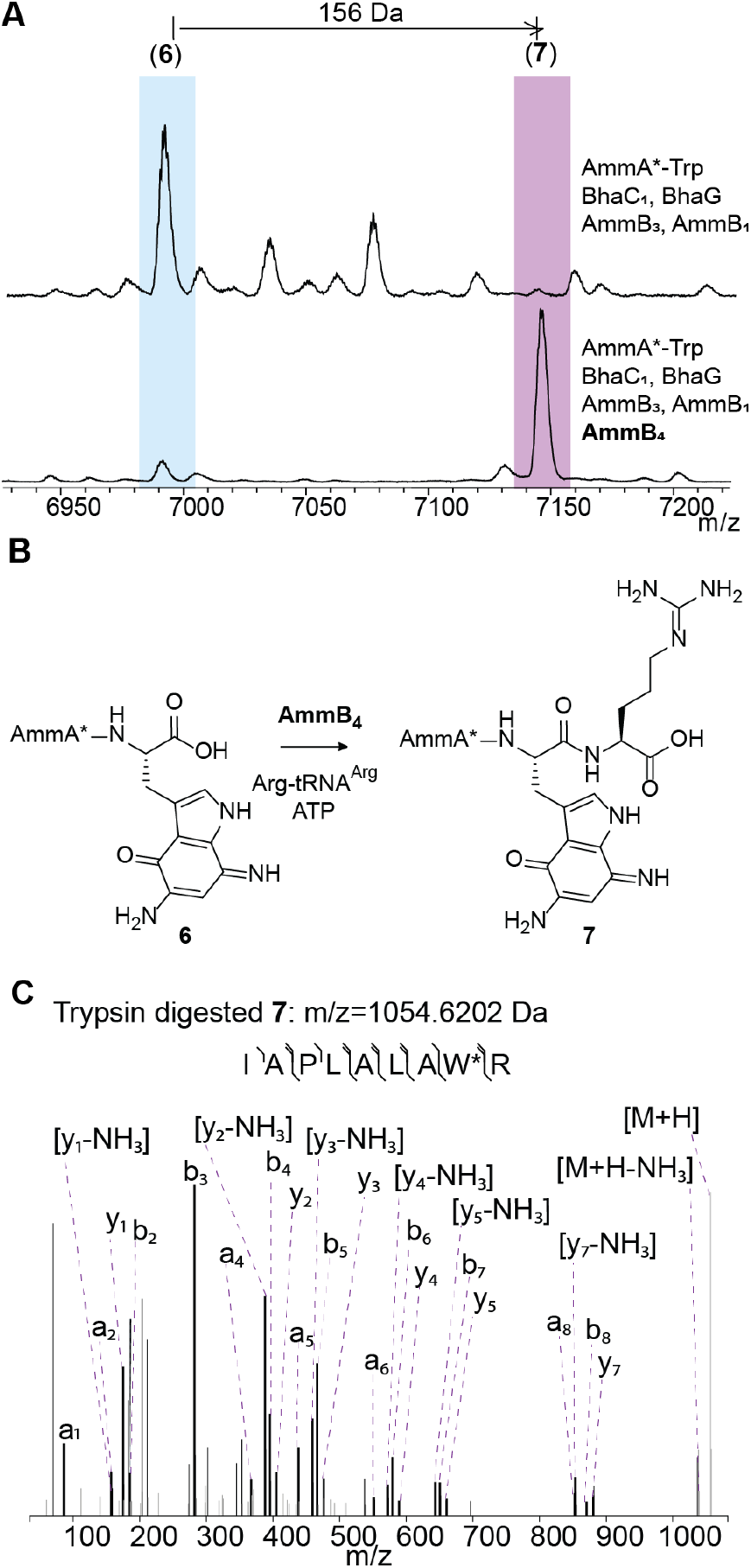
(A) MALDI-TOF mass spectra of the products of coexpression of His-tagged AmmA*-Trp with (top) BhaC_1_ (AmmC_1_ ortholog),^16^ BhaG (Amm14 ortholog),^16^ AmmB_3_, and AmmB_1_, and (bottom) additional co-expression with AmmB_4_. Calculated average m/z for **6**: 6986.1 Da, observed average m/z for **6**: 6986.4 Da. Calculated average m/z for **7**: 7142.2 Da, observed average m/z for **7**: 7142.3 Da. (B) Reaction scheme of the new product formed during in vitro reaction of purified intermediate **6** in the presence of AmmB_4_, Arg-tRNA^Arg^, and ATP. (C) MS/MS analysis of a trypsin digest fragment of **7** supporting the addition of Arg to the C-terminus of intermediate **6**. Calculated monoisotopic m/z for peptide shown: 1054.6156 Da. See Table S2 for the m/z data of the fragment ions.

PEARLs like AmmB_4_ have been shown to add amino acids in an aminoacyl-tRNA and ATP dependent manner to the C-termini of peptides including Ala, Trp, Cys, Gly, Asn, Leu, and Thr.^15-18^ ATP-dependent phosphorylation of the C-terminus of the peptide substrate is required for carboxylate activation,^19^ and recent structure-based analysis revealed that these enzymes may have evolved from the family of ATP-GRASP enzymes.^16^ Nucleophilic attack from the amine group of an aminoacylated tRNA then forms a new amide bond, and hydrolysis of the tRNA liberates the scaffold peptide extended by a single amino acid. To confirm that AmmB_4_ catalyzed the appendage of Arg in an arginyl-tRNA^Arg^-dependent manner, His-tagged AmmB_4_, *E. coli* ArgRS, and in vitro transcribed tRNA^Arg^ were all prepared and purified according to previously reported protocols.^16^ In vitro reactions with the purified diaminoquinone intermediate **6** were performed using unlabeled Arg and L-^13^C_6_-Arg. As expected, the addition of unlabeled and labeled Arg was observed by MALDI-TOF MS with mass shifts of +156 and +162 Da, respectively (Figure S3). Furthermore, the incorporation of the isotopically labeled Arg was confirmed with high-resolution MS fragmentation analysis as shown in Figure S4. Thus, AmmB_4_ is the first PEARL to use a charged amino acid linked to tRNA, which will aid future studies on the factors that determine the specificity of these enzymes and potentially allow prediction of their substrates.

To investigate if structure prediction could provide insight into how AmmB_4_ recognizes an amino acid with a charged side chain, an AlphaFold3 model^20^ was made of the complex of the enzyme, tRNA^Arg^, ATP and the peptide AmmA*W to mimic intermediate **6**. As observed previously,^21^ the model placed the 3’-CCA sequence of the tRNA right next to the C-terminus of the AmmA*W peptide and the γ-phosphate of ATP (Figure S5). An electrostatic surface potential map of the enzyme showed that in addition to the positively charged patches where the acceptor stem and the anticodon loop of the tRNA as well as ATP interact with the enzyme, a negatively charged area is seen arising from Glu581 that we tentatively assign as interacting with the guanidinium group of the Arg in Arg-tRNA (Figure S5). Glu581 is absent in all previously characterized PEARLs.

### Heterodimeric protease Amm12/Amm13 cleaves intermediate 7

After several attempts to co-express intermediate **7** with additional enzymes in the BGC including Amm3 (the predicted halogenase)^10^ without observing any further modifications (Figure S6), we hypothesized that the next step in the pathway could involve C-terminal proteolysis to release a shorter peptide containing the aminoquinone scaffold. A search using HHpred indicated that one of the enzymes encoded in the BGC, Amm19, belongs to the M20 peptidase family with homology (37% sequence identity) to a member of a M20C subfamily of Xaa-Arg dipeptidases. Amm19 was expressed using an *E. coli* codon-optimized gene as an N-terminally His-tagged protein and purified. Incubation of His-Amm19 with peptide **7** did not produce any new products when analyzed by MALDI-TOF MS (Fig. 3B).We next turned to Amm12 and Amm13, which display homology to the heterodimeric metalloprotease TldD/TldE.^22,23^ An AlphaFold3 predicted structure of heterodimeric Amm12/Amm13 (Figure S7) showed a similar fold and dimer interaction as observed in the TldD/TldE crystal structure (PDB 5NJC).^23^ To test the involvement of these proteases in the next biosynthetic step, we expressed in *E. coli* N-terminally His-tagged Amm12 by inserting its *E. coli* codon-optimized gene into the first multiple cloning site (MCS) of a pRSFDuet-1 vector. We co-expressed untagged Amm13 with its gene inserted into the second MCS. During Ni^2+^ affinity purification two bands were observed indicating that Amm13 was successfully pulled down with His-tagged Amm12 (Fig. S8). Incubation of the purified arginine-containing intermediate **7** and purified Amm12/Amm13 yielded a new ion observed by MALDI-TOF MS with a mass decreased by 386 Da compared to **7** (Figure 3B), consistent with the C-terminal removal of a dipeptide by proteolysis. To confirm the formation of the dipeptide (expected m/z 405.1993 Da), the reaction mixture was desalted and analyzed by high-resolution electrospray ionization (ESI) MS-MS, which confirmed the formation of dipeptide **8** (Figure 3C). To corroborate the requirement for a heterodimeric protease, we also expressed and purified His-tagged Amm12 and Amm13 individually. Incubation of peptide **7** with purified Amm12 or Amm13 in separate reactions did not result in any cleavage (Figure S9).

**Figure 3.**
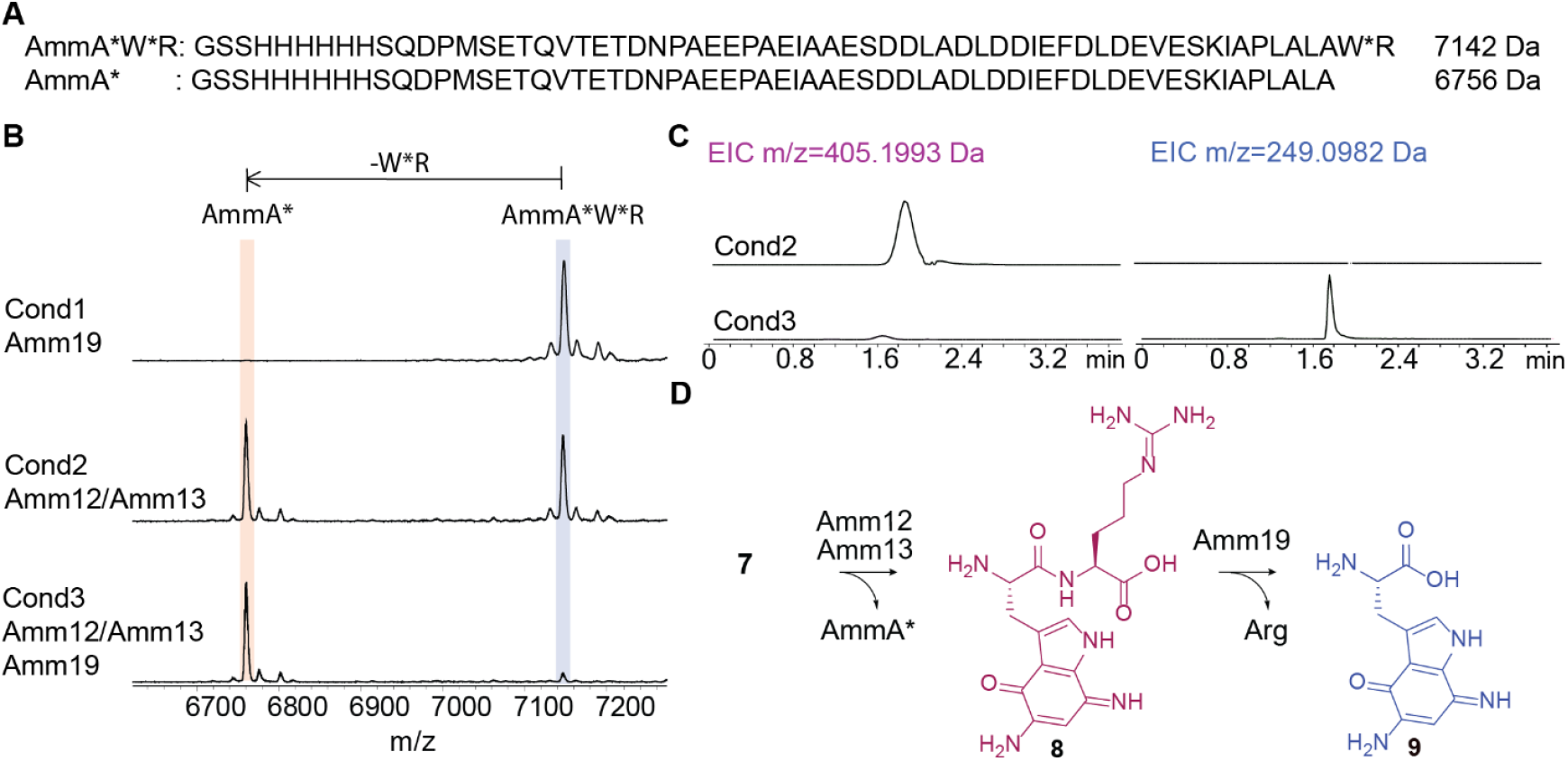
(A) Amino acid sequence for intermediate **7** in comparison with the peptide sequence of AmmA*.(B) MALDI-TOF MS analysis of the incubation of intermediate **7** with the two proteases found in the *amm* BGC, Amm19 and Amm12/Amm13. Calculated average m/z for **7**, 7142.2 Da, observed average m/z for **7**, 7142.3 Da. Calculated average m/z for AmmA*, 6756.0 Da, observed average m/z for AmmA*, 6756.1 Da. (C) LC-MS analysis of the reaction mixtures obtained from incubation with Amm12/Amm13 (cond2) versus incubation with Amm12/Amm13 and Amm19 (cond3). (D) Reaction scheme of the proteolytic cleavage catalyzed by Amm12/Amm13 followed by dipeptidase activity by Amm19.

To evaluate the requirement for an Arg in intermediate **7** for proteolytic activity by Amm12/13, we incubated every peptide intermediate preceding **7** in the pathway with purified Amm12/Amm13. Analysis by MALDI-TOF MS showed that none of these intermediates (**1-6**) were processed by Amm12/Amm13 (Figure S10). We next analyzed whether an analog of intermediate **7** containing an Arg but without any Trp modification would be recognized by Amm12/Amm13. We expressed the peptide sequence AmmA*WR and showed that it was processed by Amm12/Amm13 as observed by the cleavage of the C-terminal dipeptide by MALDI-TOF and MS/MS analysis (Figure S11). We also investigated the specificity for a C-terminal Arg by modifying the C-terminal amino acid to phenylalanine, alanine, lysine, and glutamic acid. We observed a preference for C-terminal arginine compared to these other peptides, which all cleaved much less efficiently. Extended incubation showed that the peptide variants were ultimately processed except the analog with a C-terminal glutamic acid, alanine, and lysine (Figure S11). Thus, Amm12/Amm13 appears to recognize the C-terminal sequence of **7** for specific removal of the last two amino acids as a dipeptide whereas shorter peptides that are structurally similar are not accepted. Thus, these specificity experiments show that Amm12/13 will only act after AmmB_4_ has added the Arg residue. This specificity is very different from the other well characterized TldD/E enzyme involved in the maturation of another RiPP, microcin B17, which removes a twenty-six-amino acid leader peptide in a non-sequence specific step-wise manner.^23^ Other TldD/E enzymes are more similar to Amm12/13 in that they have been shown to be endopeptidases that remove leader peptides in one step.^24-26^

### Amm19 cleaves the dipeptide 8 releasing modified Trp

After confirming the activity of Amm12/Amm13, the purified intermediate **7** was incubated with purified Amm12/Amm13 and Amm19, and the reaction was analyzed by MALDI-TOF MS. When both proteases were added to the reaction mixture, the peak of AmmA* was again observed (Figure 3B), indicating that Amm19 did not further process this peptide. However, when the same reaction mixture was analyzed by HR ESI-MS/MS, the dipeptide of 405 Da was no longer present, and the appearance of a new mass of 249 Da (**9**) was observed (Figure 3C). This observation indicated that the putative dipeptidase Amm19 cleaves the dipeptide produced by Amm12/Amm13, thereby liberating the modified Trp.

To further establish the chemical nature of the modified Trp, the reaction was scaled up and the sample was purified by HPLC and analyzed by 2D NMR spectroscopy. The corresponding signals indicated the presence of the α and β protons of Trp and the indole protons at the C-2 and C-6 positions (Figure S12). Based on the observed mass and NMR data, the product was assigned structure **9** (Figure 3D).

### The putative chlorinase Amm3 catalyzes the chlorination of 9

Phylogenetic analysis in a previous study^10^ indicated that Amm3 clustered with MibH, a halogenase that postranslationally installs a chlorine onto the side chain of a Trp in a peptide during biosynthesis of the lantibiotic NAI-107.^27^ Accordingly, Amm3 was predicted to be a flavin-dependent enzyme that would require FADH_2_ for catalysis and act on a peptide, but as noted previously, peptidic substrates **2-7** were not substrates for Amm3. Initial studies on His-Amm3 expressed in *E. coli* using a codon-optimized gene indicated that the enzyme was challenging to purify and did not contain any flavin cofactor as observed by the lack of the typical yellow color. Therefore, our first goal was to find conditions to purify Amm3 with the flavin cofactor bound. Previous studies on halogenases demonstrated the benefit of using chaperones.^27,28^ We therefore co-expressed His-tagged Amm3 with a plasmid encoding untagged chaperones GroES/EL. After Ni^2+^-affinity purification as indicated in the Methods section, the eluted His-tagged protein was yellow indicating the presence of flavin. An aliquot containing the purified Amm3 was boiled to release the cofactor, which was identified as FAD using MALDI-TOF MS (Figure S13). With the purified His_6_-Amm3 in hand, we tested intermediate **9** as a potential substrate using FAD, an NADH-dependent FAD reductase (MibS),^27^ and NADH (Figure S13) without observing any chlorination; HR MS/MS analysis only detected the cyclized pyrroloiminoquinone core **10a** that spontaneously formed under the reaction conditions.

A DALI search using an AlphaFold3-predicted structure of Amm3 identified BorH, a chlorinase involved in the biosynthesis of the bisindole alkaloid borregomycin A, as a structural homolog. During in vitro studies, BorH was shown to be inhibited by excess FAD.^29,30^ Therefore, we attempted the halogenation reaction without adding FAD. Under these conditions, we observed complete disappearance of the peak for **9** and the appearance of a new peak corresponding to the chlorinated cyclized intermediate **10b** as shown in Figure 4B. This intermediate (ammosamaic acid) was previously observed after genetic deletions of *amm4*, a putative F420-dependent oxidoreductase, as well as deletion of *amm3* with genetic complementation of *amm3*.^10^ In the in vitro chlorination reaction of **9** with Amm3, we observed product **10b**, but also cyclized non-chlorinated **10a** whereas **9** was completely consumed. We conclude from this observation that **9** is the substrate for chlorination and not **10a**. We also performed the reaction in the presence of NaBr, which resulted in the incorporation of bromine into the pyrroloiminoquinone scaffold, forming the corresponding brominated analog **10c** (Fig. 4A).

**Figure 4.**
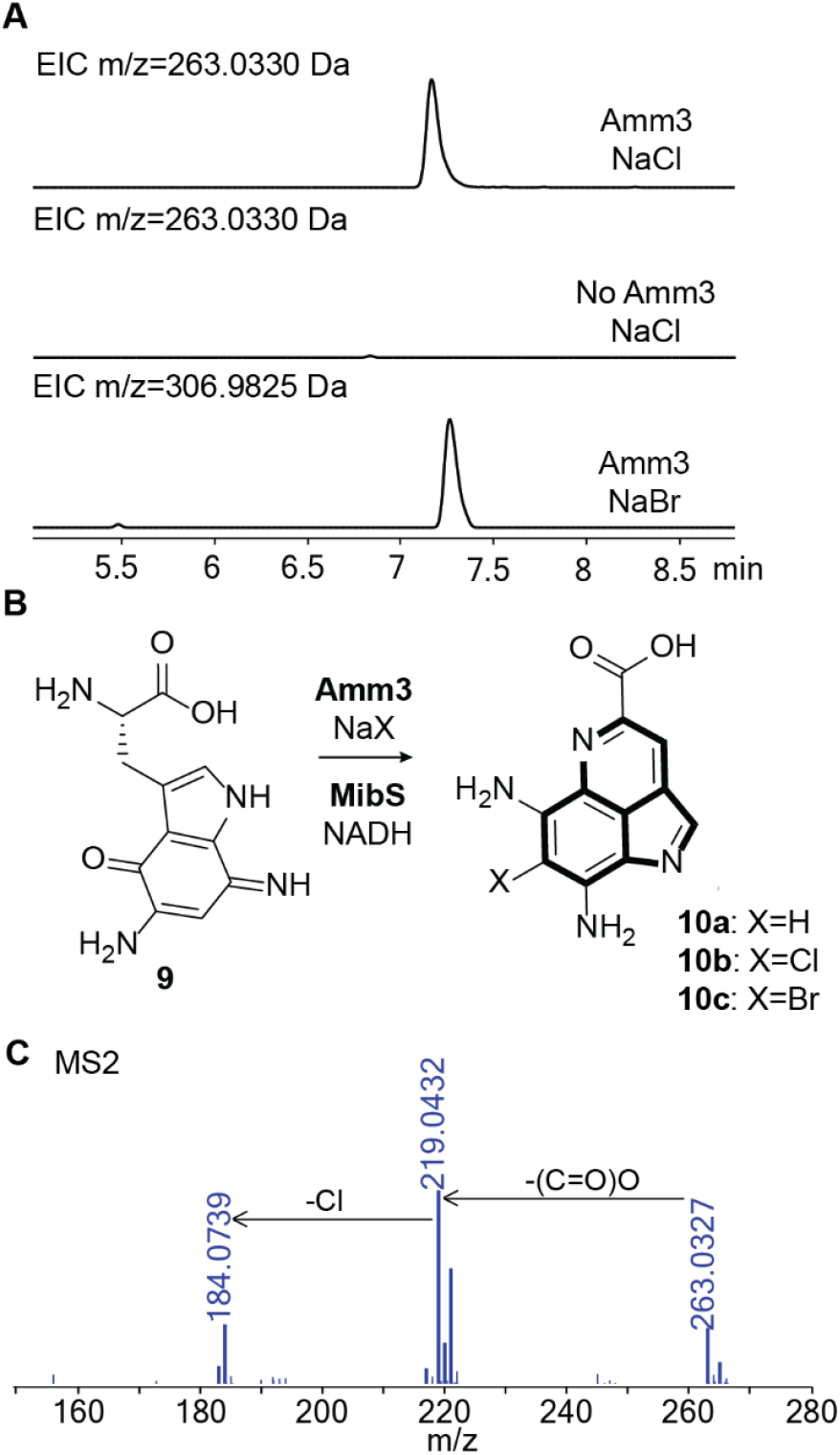
(A) Extracted Ion Chromatogram (EIC) traces of the reactions with HPLC-purified **9** in the presence of Amm3 and NaCl (top), in the absence of Amm3 with NaCl (middle), and in the presence of Amm3 and NaBr (bottom). All reactions contained MibS and NADH. (B) Reaction scheme of the new product formed during incubation of **9** with Amm3 and the corresponding NADH-dependent FAD reductase, MibS. (C) MS/MS analysis of the chlorinated product **10** showing the isotopic pattern expected for Cl incorporation. Calculated monoisotopic m/z for **10**: 263.0330.

To make sure that the conditions identified for Amm3 activity did not result in chlorination of peptidic intermediates that had been tested previously, we tested each individual peptide intermediate **2-8** with Amm3 under the new reaction conditions, however, no mass shift was observed by MALDI-TOF MS (Figure S14), indicating that the substrate for Amm3 is the highly decorated Trp derived diaminoquinone **9**. Cyclization of **9** or chlorinated **9** to the ammosamaic acid derivatives occurs spontaneously, preventing the detection of (uncyclized) chlorinated **9**. Several attempts were performed to trap chlorinated **9** by changing the enzyme load or by quenching the reaction at earlier times. However, we only detected **10a** or **10b**. With the demonstration that Amm3 acts on small molecules, the number of halogenases that act on Trp inside peptides remains confined to the chlorinase MibH,^27^ the brominases SrpI^31,32^ and MppI,^33^ and the substrate tolerant enzyme ChlH from a lasso peptide BGC.^34^

We briefly tested the substrate scope of Amm3 with several Trp derivatives as well as naphthalene-derived compounds (Figure S15 and S16). Trp was converted to 6-Cl-Trp by Amm3 as shown by NMR analysis and comparison with a commercially available reference, albeit without complete conversion. Upon scale up, we were able to also detect a second minor product of the reaction as 5-Cl-Trp. Amm3 also accepted 5-amino-indole and converted it to three different constitutional isomers of the chlorinated product. However the naphthalene derivatives were not substrates for Amm3.

### The putative asparagine synthase Amm20 may amidate intermediate 9

We hypothesized that a putative amidotransferase, Amm20, in the *amm* BGC could catalyze one of the last required modifications that converts the carboxylic acid at the C4 position of the pyrroloquinoline core into an amide group. In support of this hypothesis, in an orthologous BGC from *Streptomyces uncialis* DCA2648, an organism that does not produce an amidated pyrroloquinoline core but instead ammosamide esters, the gene for Amm20, annotated as asparagine synthase, is absent.^35^ An alternative early proposal that Amm4, a predicted F420-dependant oxidoreductase, could be involved in this step is less likely after the *S. uncialis* BGC was shown to contain Amm4 yet this organism does not make ammosamide C.^35^ To test the involvement of Amm20 in the amidation step, we constructed a C-terminal His-tagged construct using an *E. coli* codon-optimized gene (the N-terminus is required for catalysis^36^), but under all conditions tried (different temperatures, induction conditions and use of chaperones) the enzyme was insoluble. Thus, we were not able to experimentally verify that Amm20 converts the carboxylate of **10a** or **10b** into the corresponding amide.

### Amm23 methylates chlorinated ammosamaic acid

Bioinformatic analysis of Amm23 suggests that it is an *S-*adenosylmethionine (SAM) dependent methyltransferase. The gene was cloned into a pRSF-Duet vector for expression with an N-terminal His-tag using an *E. coli* codon-optimized gene, and the protein was expressed and purified using Ni^2+^-affinity chromatography. In the presence of SAM, we observed methylation of the chlorinated intermediate **10b** (Figure S17), whereas no activity was observed with **9** or **10a**. Prior work identified intermediate **11** (Figure 5) in genetic knockout experiments when *amm23* was deleted. Thus, while we observed methylation activity with **10b**, we cannot rule out that the physiological substrate for Amm23 is compound **11** after the amidation by Amm20.

**Figure 5.**
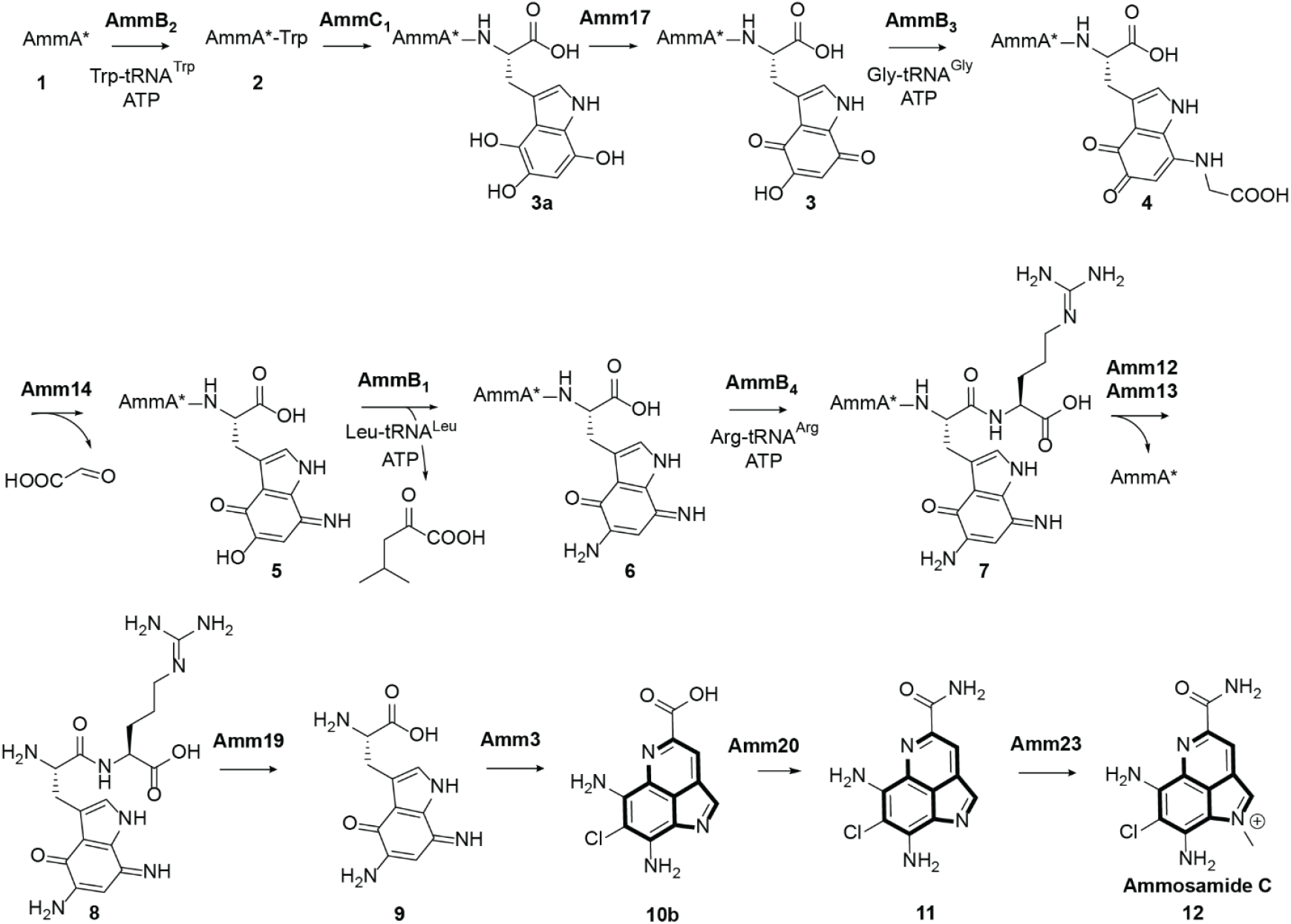
Biosynthetic steps required for the formation of ammosamide C in *Streptomyces* sp. CNR698.

## Conclusions

The biosynthetic pathway to the ammosamides has been intriguing. The BGC is very large for the formation of this group of compounds of seemingly low structural complexity. Previous studies showed that they are made by posttranslational modification involving several unprecedented transformations. In this work, we presented in vivo and in vitro characterization of the remaining steps to complete ammosamide formation that again revealed several surprises. First, the last previously uncharacterized PEARL (AmmB_4_) encoded in the BGC adds an Arg to the C-terminus of **7**. Interestingly, unlike all previous pathways involving PEARLs, in the case of AmmB_4_ no atoms of the newly added C-terminal Arg end up in the final product. A possible explanation for this seemingly unnecessary step in the pathway could have been that the amino group of the Arg serves as the donor for the amide in ammosamide A, but such a role is not supported by any of our data. Instead, the addition of the Arg appears to ensure recognition by a TldD/E analog that cleaves off the C-terminal dipeptide, possibly to avoid premature release of intermediates in the pathway. Our data also show that not all steps that could have been performed on the scaffold peptide are performed on a peptidic substrate. While the cyclization step necessitates prior cleavage from the peptide, the chlorination, amidation, and methylation in principle could all have been performed on a peptide intermediate, but instead involve small molecule substrates after proteolytic release of an advanced intermediate. These observations may also have implications for other pearlins, the group of natural products made by posttranslational modifications on the end of a scaffold peptide.^37^

The data in this work show that of the fourteen genes in the putative ammosamide BGC for which the function had not yet been assigned in previous studies, seven are required to form ammosamide C. This leaves seven other genes that are still functionally uncharacterized. Some of the proteins encoded by these genes likely do not carry out chemistry such as the transporter Amm21, the tetratricopeptide repeat containing protein AmmC_2_ that may be important for protein-protein interactions in a multi-enzyme complex,^38,39^ and the cation/H^+^ exchanger transmembrane protein Amm2 (PF00999) that may or may not be part of the BGC. Of the remaining four genes, it is likely that *amm4* is involved in the chlorination step based on gene disruption experiments.^10^ Since Amm4 is predicted to be a F420-dependent enzyme, we could not test its activity in *E. coli*, which does not have the ability to produce this cofactor. Amm10 is conserved among other putative ammosamide-like BGCs including the lymphostin BGC (Figure S18) and is annotated as a hypothetical protein. Amm15 is annotated as a glyoxalase-like domain-containing protein (PF13468) and Amm16 is a predicted oxidoreductase (PF02441). Analysis of homologous BGCs including those that produce lymphostin and ammosamide esters (Figure S18) indicates that *amm15* and *amm16* are likely not required for production of the common intermediate deschloro ammosamaic acid, **9**, because these genes are absent in these BGCs. The biosynthetic steps in Figure 5 require several interconversions between quinone and hydroquinone forms of the intermediate peptides,^16^ and it is possible that *amm16* (like *amm17*)^16^ is involved in such steps but that other enzymes can substitute.

Because Amm10 did not show any similarity to known enzyme families, a structural homology search was performed using an AlphaFold3 model of Amm10 and submitted to the Dali Server^40^ as well as to HHPRED.^40,41^ This search resulted in retrieval of proteins with only partial homology and with low homology scores. Amm10 appears to contain a C-terminus highly similar to a RiPP recognition element (RRE)^42^ whereas the N-terminus shows similarity to specific regions of TsrM, a radical SAM methylase (Z score 7.7 and 15% sequence identity), and AprD4, a radical SAM dependent enzyme that catalyzes the 1,2-diol dehydration of paromamine in the biosynthesis of apramycin (Z score 7.6 and 17% sequence identity). But Amm10 does not contain the typical cysteine motif characteristic of radical SAM dependent enzymes. Hence the role that Amm10 might play remains unclear.

In summary, this study reveals an unexpectedly lengthy route to ammosamide C that illustrates the opportunistic nature of natural product biosynthesis in which enzymes derived from newly recruited genes drive pathways that are not necessarily the most direct chemical routes to the final product. This work also demonstrates the use for a PEARL that is unlike previous roles, identifies steps that are not PTMs, and adds Arg-tRNA to the growing repertoire of amino acyl tRNAs that are used by PEARLs.

## ASSOCIATED CONTENT

### Supporting Information

The Supporting Information is available free of charge at https://pubs.acs.org/doi/

Experimental procedures, Figures S1-S18 showing assay data and AlphaFold models, and Tables S1-S2 with nucleotide sequences, primers, and MS data.

## AUTHOR INFORMATION

### Authors

Josseline S. Ramos Figueroa – Department of Chemistry and Howard Hughes Medical Institute, University of Illinois at Urbana-Champaign, Urbana, IL, United States.

Matthew Halliman, Department of Chemistry and Howard Hughes Medical Institute, University of Illinois at Urbana-Champaign, Urbana, IL, United States.

### Notes

The authors declare no competing financial interests

