## Supplemental Information for "The Biosynthetic Pathway to the Pyrroloiminoquinone Marine Natural Product Ammosamide C"

All MS and NMR data are deposited at:

Ramos Figueroa, Josseline; van der Donk, Wilfred (2025), “Data associated with "The Biosynthetic Pathway to the Pyrroloiminoquinone Marine Natural Product Ammosamide C"”, Mendeley Data, V1, doi: 10.17632/7hj55cr9zx.1

Data will be release upon publication.

### Materials and Methods

Plasmid construction and maintenance and protein overexpression were conducted using *Escherichia coli* DH10β and BL21 (DE3) strains, respectively. DNA sequences for primers and gBlocks were ordered from Integrated DNA Technologies (see Table S1). Q5 HF DNA Polymerase and deoxynucleoside triphosphate (dNTP) was purchased from NEB. NEBuilder HiFi DNA Assembly master mix was used for plasmid cloning following Gibson assembly protocols. Super DHB used as matrix in matrix-assisted laser desorption/ionization time-of-flight (MALDI-TOF) mass spectrometry (MS) was purchased from Sigma-Aldrich. $C_{18}$ resin ziptip pipette tips were purchased from Millipore Sigma. The NuPAGE™ 4%-12%, Bis-Tris, precast polyacrylamide gels, NuPAGE™ LDS sample buffer (4X), NuPAGE™ MES SDS running buffer (20X), and NuPAGE™ sample reducing agent (X10) used for SDS-PAGE were purchased from Thermo Fisher. Precision Plus Protein™ All Blue Protein ladder was purchased from Bio-Rad. Molecular grade water, and RNase/DNase free low-binding tubes were purchased from Fisher Scientific. DNA miniprep kits were purchased from Qiagen. Polymerase chain reaction (PCR) was performed using a Bio-Rad C1000 thermocycler. MALDI-TOF MS analysis was carried out in the mass spectrometry facility at the University of Illinois at Urbana-Champaign using a Bruker Daltonics UltrafleXtreme MALDI-TOF mass spectrometer. MALDI-TOF MS data were calibrated using Protein Calibration Standard I purchased from Bruker and processed using the software FlexAnalysis. Super-DHB matrix with a stock concentration of 25 mg/mL was mixed with samples in a 1 to 1 ratio and air-dried before MALDI-TOF-MS analysis. Structural predictions were performed using the AlphaFold3 server. Tandem mass spectrometry MS1 and MS2 data were obtained using an Agilent qTOF instrument equipped with an UPLC system. Two-dimensional (2D) nuclear magnetic resonance (NMR) experiments were recorded using a Bruker 500 MHz Avance III HD or 600 MHz Avance NEO spectrometer, both equipped with a 5 mm cryoprobe at 25 $^{o}$C. The samples were in 90% $H_2O$ and 10%$D_2O$. Ni-NTA resin for peptide and protein purifications was obtained from Thermo Fisher.

### Peptide and protein expression and purification

Posttranslationally modified peptides were expressed by growing cultures of *E. coli* BL21 (DE3) cells co-transformed with plasmids bearing genes encoding N-terminal His-tagged AmmA*W (see

Figure S1) and modifying enzyme. pACYCDuet-1 was used to clone the gene sequences for His-tagged AmmA* in the first multiple cloning site (MCS); untagged PEARL sequences corresponding to $AmmB_3$ and $AmmB_1$ were cloned in the second MCS. $AmmB_3$ and $AmmB_1$ gene sequences were separated by a ribosomal binding site sequence (aaggagatataca). pCDFDuet-1 was used from a previously reported study[1] encoding an $AmmC_1$ homolog ($BhaC_1$) and Amm14 homolog (BhaG). For in vitro enzyme assays, genes encoding $AmmB_4$, Amm12, Amm13, Amm19, Amm20, Amm23, and *E. coli* ArgRS were cloned as His-tagged fusion constructs using a pET28b(+) vector. Plasmids were individually used to transform *E. coli* BL21 (DE3) electrocompetent cells. pET28b(+) harboring Amm3 was co-expressed with chaperones GroEL/ES expressed from a pACYC-Duet1 plasmid.[2,3] Single colonies were used to inoculate LB cultures with the corresponding antibiotics and grown overnight at 37 °C and shaken at 200 RPM. Overnight cultures were used to seed larger cultures that were grown until the OD600 reached 0.6-0.8 at which time expression was induced with IPTG to 0.4 mM final concentration. The cultures were then grown overnight at 18 °C and shaken at 200 RPM. Peptide expression to obtain modified peptide required up to 80 L of LB to obtain sufficient amounts for structural characterization. Cells were harvested in a centrifugal device at 5,200 ×*g* and the pellets were stored at −80 °C for future use or immediately resuspended in denaturing (6 M guanidinium hydrochloride, 50 mM HEPES, 300 mM NaCl, and 50 mM imidazole at pH 7.5; for peptides that were expressed insolubly) or native buffer (50 mM HEPES, 300 mM NaCl, and 50 mM imidazole, 5% glycerol at pH 7.5; for enzymes) containing Pierce protein inhibitor cocktail and 1 mM TCEP.

<u>Peptide purification:</u> Posttranslationally modified peptides were purified from cell paste resuspensions using denaturing buffer. All peptides were processed by sonication over ice with cycles of 2 s on and 5 s off for 5 min total and an amplitude of 50%. After cell lysis, centrifugation was performed at 50,000 ×*g* for 30 min and the supernatant was loaded onto a Ni-NTA resin previously equilibrated with start buffer (6 M guanidinium hydrochloride, 50 mM HEPES, 300 mM NaCl, and 50 mM imidazole at pH 7.5). After washing with 20 column volumes (CV) of wash buffer containing 50 mM HEPES, 300 mM NaCl, 50 mM imidazole, at pH 7.5 to remove non-specifically bound protein, the His-tagged peptide was eluted with 5 CV of 50 mM HEPES, 300 mM NaCl, 200 mM imidazole, at pH 7.5. The eluting solution was then used for HPLC purification or desalted using C18 ziptip and eluted in a 1.5 µL solution of Super-DHB of 25 mg/mL

concentration (dissolved in 80% $CH_3CN$ (ACN)/20% $H_2O$/0.1% $CF_3COOH$ (TFA)) for MALDI-TOF MS analysis.

Protein purification: All other proteins mentioned above were purified individually after resuspension of the cell pellet in 50 mM HEPES, 300 mM NaCl, 50 mM imidazole, 5% glycerol at pH 7.5. As for the peptide purification, lysis was performed by sonication over ice with 2 s on and 5 s off for total of 5 min on and amplitude of 40% magnitude. After centrifugation at 50,000 ×*g* for 30 min, the supernatant was loaded onto a Ni-NTA resin previously equilibrated with 50 mM HEPES, 300 mM NaCl, 50 mM imidazole, 5% glycerol at pH 7.5. After washing with 20 CV of 50 mM HEPES, 300 mM NaCl, 50 mM imidazole, 5% glycerol at pH 7.5 to remove non-specifically bound proteins, the His-tagged protein was eluted with 5 CV of 50 mM HEPES, 300 mM NaCl, 200 mM imidazole, 5% glycerol at pH 7.5. The excess imidazole content was then removed by concentration and buffer exchange into 50 mM HEPES, 300 mM NaCl, 5% glycerol at pH 7.5 using the appropriate MWCO Amicon centrifugal device. Amm3 purification followed the same protocol, but different start, wash, and elution buffers were used. Start and wash buffer contained 20 mM $NaH_2PO_4$, 500 mM NaCl, and 50 mM imidazole at pH 7.5 with 5% glycerol, and elution buffer consisted of 20 mM $NaH_2PO_4$, 500 mM NaCl, and 200 mM imidazole at pH 7.5 with 5% glycerol, while storage buffer (after imidazole removal) contained 5.0 mM $K_3PO_4$ and 5% glycerol at pH 8.0.

**High-performance liquid chromatography (HPLC) purification of 6 to 9**

Peptides obtained from Ni purification were acidified with a 10% trifluoroacetic acid (TFA) solution to a final concentration of 0.1 to 2% TFA to precipitate any coeluting protein from the overexpression protocol. The sample was then passed through a 0.45 μm syringe filter and injected directly into an Agilent 1260 Infinity II HPLC system for purification. The peptide was purified using a preparative scale column, VP NUCLEODUR C18 HTec, 5 μm, 250 x 10 mm with the following gradient: 0-5 min isocratic 2% acetonitrile (ACN), 5-40 min 2%-60% ACN, 40-42 min 60%-100% ACN; mobile phase contained 0.1% TFA as reported in previous work.[1] Fractions were collected manually and were submitted for MALDI-TOF MS analysis by combining 1 to 1 with a 25 mg/mL solution of SDHB (prepared as mentioned above). Fractions containing the desired peptide (usually bright purple in color) were combined and lyophilized. The dried purple solids were then used as substrates for *in vitro* studies, further purified on an analytical scale column for structure elucidation, or digested for LC-MS or NMR analysis when required.

When an additional round of purification was required, peptides were redissolved in deionized water, passed through a 0.45 µm filter, and injected directly onto an analytical scale Agilent 1260 Infinity HPLC system. A Vydac 218TP C18 column, 5 µm, 250 x 4.6 mm was used for peptide purification using the following gradient: 0-10 min isocratic 2% ACN, 10-30 min 2%-60% ACN, 30-32 min 60%-100% ACN; mobile phase contained 0.1% TFA. Desired peptide was identified by MALDI-TOF MS analysis performed on each collected fraction.

Dipeptide **8** and compound **9** required a different column for purification. After the *in vitro* reaction of peptide **7** with Amm12/Amm13, and with Amm12/Amm13 and Amm19, respectively, dipeptide **8** and compound **9** were purified by a first round of purification using a Vydac 218TP C18 column using the following gradient: 0-10 min isocratic 2% ACN, 10-30 min 2%-60% ACN, 30-32 min 60%-100% ACN; mobile phase contained 0.1% TFA. After detecting an impurity in the purified peptide by NMR spectroscopy, a second round of purification was performed by using a Chromosorb Varian C18 column using the following gradient: 0-10 min isocratic 2% ACN, 10-30 min 2%-60% ACN, 30-32 min 60%-100% ACN; mobile phase buffered with 0.1% TFA.

When the samples were used for NMR experiments or LC-MS/MS analysis, HPLC-purified **7** to **9** were first subjected to trypsin or chymotrypsin proteolytic digest in a 1 to 20 w/w ratio. The samples were then acidified to 0.1 to 1% TFA, filtered through 0.45 µm membrane or centrifuged at 13,000 RPM for 5 min, and injected onto the HPLC for further purification. The fragment containing the C-terminal modifications were then eluted with the same column and gradient as indicated above for the analytical scale purification. Fractions were collected manually and submitted for high resolution (HR) MS analysis for peptide fragment identification and NMR analysis.

**LC-MS analysis of compounds 7 to 10**

After trypsin digestion (for **7**) or after *in vitro* reactions performed to obtain **8** to **10**, each solution was desalted using a Pierce™ C18 spin column. Briefly, the spin column was activated with 200 µL of 60% ACN with 0.1% formic acid (FA) and rinsed three times with 200 µL of 0.1% FA. The sample was loaded onto the equilibrated column and the column was washed three times with 200 µL of 0.1% FA. Then, the compounds of interest were eluted using 25 µL of 60% ACN with 0.1% FA, twice. The elutant was lyophilized, redissolved in 25-50 µL of water, and passed through a 0.45 µm membrane to remove any insoluble material. The sample was then injected onto an Agilent 1260 Infinity II with a 6545 LC/Q-TOF mass spectrometer detector. For tryptic fragment **7**: A Kinetex® 2.6 µm C8 100 Å column, 150 x 2.1 mm was used following the gradient: 0-3 min isocratic 5% ACN, 3-13 min 5%-95% ACN, 13-15 min 95%-5% ACN; mobile phase containing 0.1% FA. For compounds **8** to **10**: A Poroshell EC-C18 column, was used following the gradient: 0-3 min isocratic 5% ACN, 3-13 min 5%-95% ACN, 13-15 min 95%-5% ACN; mobile phase containing 0.1% FA. A DualAJS ESI detector was utilized with the following parameters, Gas temperature of 325 °C, drying gas of 13 L/min, nebulizer of 35 psi, sheath gas temperature of 275 °C, sheath gas flow of 12 L/min, nozzle voltage of 500 V and MS-TOF fragmentor set at 175 V.

**AmmB$_4$ *in vitro* experiments**

In a 200 µL tube, a solution containing a final concentration of 50 mM HEPES, 25 mM KCl, and 15 mM $MgCl_2$ pH 7.6, 2 mM DTT, 5 mM L-arginine, or $^{13}C_6$-labeled L-arginine, 6 mM ATP, 10 U thermostable inorganic pyrophosphatase (TIPP), 10 U superase RNAse inhibitor, 3 ng/µL of tRNA (*E. coli* $tRNA^{Arg}$), 20 µM of the corresponding aminoacyl-tRNA synthetase ArgRS, 5 µM of PEARL AmmB$_4$, and 15 µM of peptide substrate (intermediate **6**) was incubated for 4 h at 37 °C in a heating block. After this time, the modified His-tagged peptide was isolated using Ni agarose resin. The eluted His-tagged peptide was buffer exchanged into 50 mM HEPES pH 7.5 and further cleaved with trypsin or chymotrypsin for ESI-MS/MS analysis.

**Amm12/Amm13 and Amm19 *in vitro* experiments**

HPLC-purified **7** was incubated with Amm12/Amm13 and Amm19 separately to a final protein concentration of 25 µM at 37 °C for 4 h. After this time, the solutions were desalted using C18 ziptip and eluted in a 1.5 µL solution of Super-DHB of 25 mg/mL concentration for MALDI-TOF MS analysis. Additionally, **7** was also incubated with Amm12/Amm13 and Amm19 together and after incubation for 4 h, the two reaction mixtures were desalted for MALDI-TOF analysis and HR MS/MS using Pierce™ C18 spin column.

The same procedure was repeated for intermediates in the pathway preceding **7** and the reaction mixtures were analyzed on a MALDI-TOF spectrometer.

**Amm3 *in vitro* experiments**

HPLC-purified **9** was incubated in a mixture containing 5 μM FAD-containing Amm3, 10 μM MibS, 50 mM NaCl and 2 mM NADH in a 50 mM HEPES buffer at pH 8.0 overnight in a shaker at 37 °C and shaking at 200 RPM. After this time, the solutions were desalted using a Pierce™ C18 spin column for HR MS/MS analysis. Similarly, the reaction was repeated in a mixture containing NaBr and analyzed by HR MS/MS.

To test the substrate scope of Amm3, several tryptophan (Trp) analogs (Figures S16 and S17) were evaluated as substrates for Amm3. L-Trp was shown to be a substrate and thus the chlorination site was studied. Scaled-up 50-μL reactions containing 1 mM Trp were set up ten times and an FAD-recycling system was used in an attempt to push the reaction towards higher yields of the chlorinated Trp after two-day incubation. The reaction mixture contained 25 mM HEPES, 50 mM NaCl, 1 mM NAD+, 5 mM phosphite, 1 mM Trp or Trp analog, 20 μM Amm3, 25 μM MibS, 30 μM phosphite dehydrogenase (PTDH),[4] and 0.2 μg/μL catalase. After the incubation time, the samples were processed in two ways. For Trp analogs that were not retained by Pierce™ C18 spin columns, protein was precipitated by adding an equal volume of ice cold ACN and centrifuged at 13,000 RPM for 5 min. The supernatant was then analyzed by HR MS/MS. In all other cases, desalting was performed using a Pierce™ C18 spin column and the flowthrough analyzed by HR MS/MS.

For the reaction mixtures containing Trp as a substrate, after scale-up, the reaction was acidified with TFA to a final concentration of 2%. After centrifugation at 13,000 RPM for 5 min, the supernatant was purified using an analytical scale HPLC as described above.

**Amm20 *in vitro* experiments**

HPLC-purified **9** and **10** were incubated in a mixture containing 20 μM Amm20, 50 mM L-glutamine, 2 mM ATP, and 1 mM $MgCl_2$ in 50 mM HEPES buffer containing 150 mM NaCl at pH 8.0 overnight at 37 °C. After this time, the solutions were desalted using a Pierce™ C18 spin column for HR MS/MS analysis. Similarly, the reaction was repeated in a mixture containing lysates of untagged Amm20 and analyzed by HR MS/MS.

#### Amm23 *in vitro* experiments

HPLC-purified **9** and **10** were incubated with Amm23 separately to a final protein concentration of 25 µM and 20 mM SAM in 50 mM HEPES buffer with 150 mM NaCl at pH 8.0 at 37 °C for 4 h. After this time, the solutions were desalted using a Pierce™ C18 spin column for HR MS/MS analysis.

**Table S1.** List of primers and double stranded DNA fragments applied in this study.

| DNA Name | Sequence |
|---|---|
| JRF_AW_B3_B1.VF | AAGGAGATATACAatggctgccacctcag |
| JRF_AW_B3_B1.VR | tttctttaccagactcgattagacagtttcccaacgtaa |
| JRF_AW_B3_B1.GF | gaaactgtctaatcgagtctggtaaagaaaccgc |
| JRF_AW_B3_B1-2.GR | ctgaggtggcagccatTGTATATCTCCTTttagggcgccgccac |
| JRF_gb_Amm16 | ATGACTGCACCTGTCCTTCATGCCGTTGTTACGGGAGCGCCGCCAGCCGGT<br>GATGTTGGACGCCTCGTAGCTTTGGCTGGGGCGGATGGATGGGGAACAC<br>ACCTGGTGGCATCGCCTAGTGCTCGGCCGTTCCTTGATCTTCCGAGATTAG<br>CGGCAGCAACGGGGCATCCTGTTCGGACTGATTACCGGCAACCGGGCGAT<br>CCTTGGCCTCTCCCACCACCTACCGCCGTGGTGCTGGCCCCAGCTACAGGT<br>AATACCTTAGCCAAATGGGCTGCGGGCATATCAGACACTTTAGCGCTCGG<br>TCTGCTCGTGGAAGCTGTTGGCCAGGGTACGCCAGTTGTGGCGGCACCAT<br>TCTCAAATACGGCACATCTGGCACATCCAGCGGTAGCGGAAGCAATCACT<br>CGGCTGCGCAGCTGGGGTGTTACTGTTCTCACCGGACCAGAGGTTTGTCC<br>ACCACACCCGCCGGGACCTGCAGCCGCGCATGCAGAACGTTTTGGTTGGG<br>ACGCTGTTTGGGCAGCAGTTACTGGTCATCCTCACCTGGCCGGTGCACGG<br>GCTGCCGCTCCAGCTCCGGATCCAATCCCTCAAGAGCAGCCCGCTTAG |
| JRF_gene_fragment_Amm19 | ATGTCACATAGAGATCGATGCCGGGCCGCTGTTGAACGCCGTCGTAGCAC<br>GTTATTGGAACTGTCCCACTCGCTTCACGCAGAGCCTGAGACAGCCTTTGA<br>TGAGCATCGGAGTGCGGCCAAGATTGCCGATTTACTTGAAGGTGAGGGAT<br>TTGCGGTGGAGCGTGGTACTGGCGGGTTAGCAACCGCCTTGACTGCGACA<br>TGGGGAGACGGCGAGCTCGTTGCCGGAGTTTTGGCTGAGTATGACGCCTT<br>GCCGGGGATCGGGCACGCATGCGGCCATAACGTCATAGCTGCAGCTAGT<br>GTTGGTGCGGCACTGGCCTTACGAGAGGTTGCAGCCGACCTCAACCTGAC<br>AGTCCGATTAATTGGTACTCCGGCTGAGGAATCCGGTGGCGGGAAGGTTC<br>TGCTTCTTGAGCGGGGCGCGTTTGATGATTTAGCTTTTGCGTTATTGATCC<br>ATCCGTCCCCTGACGAGATCTGCGCACCAAGAACATTGGCTGTTACCGATT<br>TGGAAGTCACTTATCGGGGCCGCGCAGCACACGCGAGCTTCGCTCCCCAC<br>CAAGGGGTAAATGCGGCAGACGCACTTACTGTAGCGCAAGTAGCTATTGG |

| | |
|---|---|
| | ATTAGCGCGTCAACATCTCGAGCCCCACCAGATGGTGCACGGGATTACTA<br>CGCACGGCGGAGACGCGCCCAACATAGTACCGGATCGAACCGGCGCGGT<br>TTATTACTTACGTGCCCGATCAGCTGAATCCCTGAGCCGGTTAGAGCATCG<br>CGTCCGGGATTGCTTACGTGCCGGTGCTGTCGCTACCGGCTGCACAGAAG<br>AGATCCGTAGAGCAGCACCGGCGTACACAGAACTGGTGTCCGATGTTCGT<br>TTGACCGCAGCATATCGAGAAGCTATTACGGCCCTCGGGCGTGTACCGCT<br>TGATGAGGAAACCGAAGCAAGACTTGTCACACCTGCTAGTACTGATATGG<br>GCAATGTTTCTCAGGTTCTTCCAGCGATTCAACCGTCGATAGCGGTCGATG<br>CCCGCGGTGCGGTTAACCATCAACCAGGCTTTGCGGAAGTTTGTGCAGGT<br>CCCTCAGGCGATGCTGCTGTCCTGGATGGTGCGGTGGCGCTTGCCTGGAC<br>CGCTGTTGCTGCTGCCAGTGATCCTGCACATCGTGTGGCATTATTAGACGG<br>CGTCCGCGCGAGACGTTGTGGCGCCGTTCGTTGA |
| JRF_gene_fragment_Ecoli_ArgRS | ATGAACATACAAGCCTTATTGTCCGAGAAAGTTCGGCAAGCTATGATAGC<br>CGCCGGTGCACCTGCGGACTGCGAACCTCAAGTGCGACAGTCCGCTAAGG<br>TGCAATTCGGGGACTACCAAGCCAACGGCATGATGGCTGTTGCTAAGAAA<br>CTGGGTATGGCTCCGCGTCAACTTGCCGAGCAGGTCTTGACACACTTGGA<br>TCTCAACGGCATCGCCTCGAAAGTGGAGATCGCTGGACCAGGCTTCATTA<br>ATATTTTCCTGGACCCAGCCTTTCTGGCCGAACATGTTCAACAAGCTTTGG<br>CTTCTGACCGGTTGGGAGTTGCAACACCGGAGAAGCAAACCATAGTAGTC<br>GATTACTCTGCACCGAACGTAGCAAAGGAGATGCATGTTGGTCACTTGCG<br>CTCGACGATCATCGGGGACGCCGCGGTTCGAACACTGGAGTTTCTTGGCC<br>ACAAAGTGATACGTGCGAACCACGTGGGCGACTGGGGTACTCAATTTGGC<br>ATGCTTATTGCGTGGTTAGAAAAGCAACAGCAGGAGAATGCTGGCGAAAT<br>GGAATTAGCCGACTTAGAGGGGTTCTACAGAGATGCGAAGAAACATTACG<br>ACGAGGACGAGGAGTTCGCCGAGCGTGCTCGCAATTACGTTGTTAAGTTG<br>CAGAGTGGTGACGAGTACTTCCGTGAGATGTGGCGGAAGCTTGTTGATAT<br>CACAATGACTCAAAACCAAATAACTTATGATCGTCTGAACGTCACGCTGAC<br>GCGGGATGACGTGATGGGTGAGAGCTTATACAACCCAATGTTGCCAGGC<br>ATAGTTGCAGACCTTAAAGCAAAGGGGCTGGCTGTGGAGAGCGAAGGTG<br>CCACTGTGGTATTCCTTGACGAGTTTAAGAATAAGGAAGGTGAGCCGATG<br>GGCGTTATCATTCAGAAGAAAGATGGCGGCTATCTTTATACAACAACCGA<br>CATTGCTTGTGCAAAGTACCGTTACGAGACCCTTCACGCCGATCGGGTTTT<br>ATATTATATCGATAGCCGTCAGCACCAACATTTGATGCAAGCATGGGCTAT<br>TGTGAGAAAAGCTGGATACGTGCCGGAATCCGTCCCGCTTGAACACCACA<br>TGTTCGGGATGATGCTCGGAAAGGACGGGAAGCCGTTTAAGACCCGAGC<br>CGGCGGAACGGTTAAACTTGCCGACCTCTTGGACGAAGCTTTAGAACGTG<br>CGCGCCGTCTGGTGGCTGAAAAGAACCCGGACATGCCGGCTGATGAACT<br>GGAGAAACTTGCGAATGCTGTTGGCATTGGGGCTGTTAAGTACGCAGACT<br>TGTCTAAGAATCGTACCACCGATTACATCTTCGATTGGGATAATATGCTTG<br>CGTTCGAGGGGAATACAGCACCTTACATGCAGTACGCTTACACTCGAGTTC<br>TTAGCGTGTTTCGCAAGGCGGAGATCGATGAGGAGCAATTGGCTGCGGC<br>GCCCGTGATTATTCGGGAAGATAGAGAGGCCCAATTGGCAGCTCGGCTGC<br>TCCAGTTCGAGGAGACGTTGACGGTAGTAGCCCGTGAGGGCACTCCACAC<br>GTTATGTGTGCGTATTTGTACGACCTCGCAGGCTTGTTCTCAGGTTTCTAC<br>GAACACTGCCCGATTCTGAGTGCTGAAAATGAGGAAGTGCGTAACTCGCG<br>ACTTAAGCTGGCCCAGTTGACTGCAAAGACGCTGAAGCTGGGACTGGATA<br>CGTTGGGGATCGAGACTGTGGAGAGAATGTAA |

| | |
|---|---|
| JRF_gene_fragment_Amm22 | ATGGCCGAGAAATCAAAACCGGGTAGCAAGCCTTACCGGAGAGACCTGCGAGAACGGTTAAAGAGCCTCGGATTTGCTGACCATCAGATCGACGACGTTGTCACAGCGGAGTTGGTACAAATGTGTCACATCCGCCCGCGTACCGCCAGAAGACTTGCAAGAGAGTTAACTTTGGACGAGGTTGCGACGCGTTACTCCAGTCTGCGTAATGATGCTGAGTCACGCATGCGCGGCTCGCGGATCTGGGACTACGAGCAATGGCCGAACCGTGGCGTGCGGCCTACAGTCCACACACTGCGTACATTGGCCGAGGTTTACGGTACAACTTGGGTAGAGCTCGTAGACATTGAGGATCTCCAACACATGCCGGAAGAAGACCGGGAACTTTACCACCTTCGGGTATCAGGCGGGCCCCCTGACACACGTGCTGCCCCTACCGCTCCCCTTCCCCCGAGTCTGCCGGCTGCACCGCCGGCCTGTGCACCTCCTCCAGCCCATGCGCACGCTCCGGCGCCAGACCGTCCTCGTCCCCGCCCAGCTCCAGCTCAACCTGACCCATCTGGTCCGGCCGGAACTGCCGAACTCATACGTACTGGTGCTGAACACGCGTTAGGAGTGGCTCGCAGCCTTAGTACCACGAATGTAGATTCCTTGACTTTACACCAATTAGAGCAAGAGTTAGCCAGTACCTCTCGGTCGTATGTTCACAGTTCCCCATATCCGTTGTATTTTCGGTTGGCACAGCTGGGCAACCACATTGCTCAATTACTGCCAGGTCGGCAGCGCCCGTCTCAAACCCGCAGACTGAATGCACTGGCCGCCCGGTGCTCTGCCTTGTTAGCGTGGGTATGCGATGATTTAGGCGCCGCAGCAGCCGCGCACGACCACGCGTGGGCAGCATGGATCTATGCATCTCACAGCGAAAGCGAGGACGCACAGCGCTGTGTAAGTCTGGTCCGTTCTCGCTTGGCATTTTCATCTGGAGACTTTATGGAGAGTGCCCGAATTGCAGATAGCGCCTGTAGTGGCGTCCCCCAAGATAGTGTCGGATCACATTTGTTGCTGCGACAAGCGTTGGCCTGGGCCCGTGCAGGGCAACACCACAAGGTACACGAGGCCCTTCGGGATTGGAAGGCAAGAGGAGAGGAAGGTCCGCTCGGAGCTGGTGACGGCATATTTGAACTTCAACAAGCGCAGCAAAGATATTTCGTAGGGGCCGCGCTGCTTGATATTTTAGACGTTGAGCATGGTCTGGACGAGCTGCGTACCGCCATCGATCTGTTCAACCGTACGCCCGAGGATCTGCGCTCATACGTGGAACACTCACTGAGCCATATTGGAACGGCGAAAGCCTTCGTGGTCCTCGGCGATGTACAAGAGGCATCCAAGGTAGTTGAGCCTGTATTCGACCTCGCACCCGAACGTCGCGTACGCGCAATCCTGTCTAGTCTCCGCGAGCTGGTCGCGTTCACGGCCGGCAGACCTTCCTACGGAAGTCAACTGGCTCGGGAGTTGTCTGAGCGAATTGACGAGTTCGCATCCCATGCTATCACTCGCCGGTTACACTCAGTTGTGTAA |
| JRF_pRSF_Amm22_VF | ATCCCATGCTATCACTCGCCGGTTACACTCAGTTGTGTAATGCAGGTCGACAAGCTTGCG |
| JRF_pRSF_Amm22_VR | GTCTCTCCGGTAAGGCTTGCTACCCGGTTTTGATTTCTCGGCCATCGGATCCTGGCTGTG |
| JRF_pRSF_Amm19_VF | GGATGGCGTACGAGCCCGCCGGTGTGGTGCAGTCCGGTAGTGCAGGTCGACAAGCTTGCG |
| JRF_pRSF_Amm19_VR | TCTTCGGCGTTCTACGGCGGCTCGGCAACGGTCGCGCATCGGATCCTGGCTGTGGTGATG |
| JRF_pRSF_Amm16_VR | CGCTCCCGTAACAACGGCATGAAGGACAGGTGCAGTCATCGGATCCTGGCTGTGGTGATG |
| JRF_pRSF_Amm16_VF | TCCAGCTCCGGATCCAATCCCTCAAGAGCAGCCCGCTTAGTGCAGGTCGACAAGCTTGCG |
| JRF_AWB3B1.F | CTGGGGTGTGTTGTTTGGCCAGCCGAAGAAGGACGCTAAtcgagtctggtaaagaaaccg |

| | |
|---|---|
| JRF_AWB3B1.R | GCGCTGTGCGTGTGGAAGTCATTGTATATCTCCTTttagacagtttcccaacgtaatgc |
| JRF_B4_AWB3B1.F | tttggcattacgttgggaaactgtctaaAAGGAGATATACAATGACTTCCACACGCACAG |
| JRF_B4_AWB3B1.R | ggcgttcaaatttcgcagcagcggtttctttaccagactcgaTTAGCGTCCTTCTTCGGC |
| JRF_pET28_A20CtHis.F | ATATTTGGCTGCGTCATTACAGTCCGGAGATCGCCGTACACCACCACCACCACCACCACC |
| JRF_pET28_A20CtHis.R | GGTATATCTCCTTCTTAAAGTTAAACAAAATTATTTCTAGAGG |
| JRF_A20_CtHis.F | CTCTAGAAATAATTTTGTTTAACTTTAAGAAGGAGATATACCATGTGTGGCATAGTCGGC |
| JRF_A20_CtHis.R | CAGCCGGATCTCAGTGGTGGTGGTGGTGGTGGTGGTGGTGGTGTACGGCGATCTCCGGAC |
| JRF_pAB1B3B4.R3 | ctcactataggggaattgtgagcggataacaattccccatcttag |
| JRF_pAB1B3B4.F | Cctgtgtttcggacatcgg |
| JRF_gDNA-A20-2F | ggcagcgtggacggcgtcagagaagggagcggacatatgATGTGCGGAATAGTCGGTTGG |
| JRF_gDNA-A20-2R | cggggatctaagcttggctgcagtcaatgatgatgatgatgatgGACCGCGATCTCCGGG |
| JRF_pET28-ArgS_R | TGGATACGTTGGGGATCGAGACTGTGGAGAGAATGTAACATATGGCTAGCATGACTGGTG |
| JRF_pET28-ArgS_F | GCCGAACTTTCTCGGACAATAAGGCTTGTATGTTCATGTGATGATGATGATGATGGCTGC |
| JRF_Ec_tRNA-Arg-ACG-r | mUmGGTGCATCCGGGAGGATTCGAACCTCCGACCGCTCGGTTCGTAGCCGAGTACTCTATCC |
| JRF_Ec_tRNA-Arg-ACG-f | AATTCCTGCAGTAATACGACTCACTATAGCATCCGTAGCTCAGCTGGATAGAGTACTCGG |
| pET28a(+)_His_Amm23 | ATGCCGGCACCTACTGACCTGCGCCTCAGAGAACTGGTAACGGGATACCGCACGTCCCAAGTCGTGTATGCCACTGTAAAACTGGGCCTGTTTGATCTGCTGGGTGCGCGCCCTCGCACGGCGGCCGAACTCGCCGAAGCCGGAGGTGCAGATCCAGATGCACTGGCACGTTTTGTACGAGCCCTGACAGCGGAAAGAATACTGCAGTCCACAGCGGACGGTGTATTATCACTGACACCACTCGGAGAAAGACTTCGTAGTGATGCACCTGGTGGGCTGGCGGCGTGGGTGGTAACGAGCTGCGAAGAGCAGTTCTTTGCATGGGGCCATGCGGTGCATACGCTGCGAACGGGCCAATCAGCATTTGAAAAAGCCCATGGAGTCGGGTTCTGGGAGCATTTACGCTCTGATGCTAAAGCGGCAGAAAGCTACGATGCAGGCATGGCCGTGTCAGCGGGTGAGGCATGCGATCTTATTGTGTCTGCCGGGGGCATCGATCGAGCTCGACATGTTGTTGATGTGGGTGGTGGCCGGGGTGCGCTCGCACGGCGGCTTCTTGATACCTTCCCACATGCGCGGGTCACCGTTCTGGACCTTCCCGAGGTCGTAGAACGTACGCGCGCGAGTTTTGCACCTGATCCAGTTCCGGAGCGCTTAGGCCTGGTACCCGGATCTTTTTTAGACGGCGTCCCTGCCGGGGGGGATGTTTACGTACTGTCACGTGTACTGGCCGATTGGGATGACGCTCATGCCGCTAGAATCCTGGCGTCTTGCCGGCGCGCCATGACACCCGGGGGGCGCCTGCTGATTGCTGAGGGACTGACTCGGGATGATGCACCCGCAGGGGCCCGCGGCTTGCTCGATTTGCATTTACTGTTGCTGTTCGGTGGACGTGAGCGCACAGTGACCGGCCTTACCGCTCTGCTGAGCGGTGCCGGATTTGCGTTGGTGTCCGTGACGGGCGCTGACAACGGAGGCATTTCGCTGCTGACGGCCGTAGCTCGGTAACTCGAGCACCACCACCACCACCACTGAGATCCGGCTGCTAAC |

| | |
|---|---|
| | AAAGCCCGAAAGGAAGCTGAGTTGGCTGCTGCCACCGCTGAGCAATAACT<br>AGCATAACCCCTTGGGGCCTCTAAACGGGTCTTGAGGGGTTTTTTGCTGAA<br>AGGAGGAACTATATCCGGATTGGCGAATGGGACGCGCCCTGTAGCGGCG<br>CATTAAGCGCGGCGGGTGTGGTGGTTACGCGCAGCGTGACCGCTACACTT<br>GCCAGCGCCCTAGCGCCCGCTCCTTTCGCTTTCTTCCCTTCCTTTCTCGCCAC<br>GTTCGCCGGCTTTCCCCGTCAAGCTCTAAATCGGGGGCTCCCTTTAGGGTT<br>CCGATTTAGTGCTTTACGGCACCTCGACCCCAAAAAACTTGATTAGGGTGA<br>TGGTTCACGTAGTGGGCCATCGCCCTGATAGACGGTTTTTCGCCCTTTGAC<br>GTTGGAGTCCACGTTCTTTAATAGTGGACTCTTGTTCCAAACTGGAACAAC<br>ACTCAACCCTATCTCGGTCTATTCTTTTGATTTATAAGGGATTTTGCCGATT<br>TCGGCCTATTGGTTAAAAAATGAGCTGATTTAACAAAAATTTAACGCGAAT<br>TTTAACAAAATATTAACGCTTACAATTTAGGTGGCACTTTTCGGGGAAATG<br>TGCGCGGAACCCCTATTTGTTTATTTTTCTAAATACATTCAAATATGTATCC<br>GCTCATGAATTAATTCTTAGAAAAACTCATCGAGCATCAAATGAAACTGCA<br>ATTTATTCATATCAGGATTATCAATACCATATTTTTGAAAAAGCCGTTTCTG<br>TAATGAAGGAGAAAACTCACCGAGGCAGTTCCATAGGATGGCAAGATCCT<br>GGTATCGGTCTGCGATTCCGACTCGTCCAACATCAATACAACCTATTAATTT<br>CCCCTCGTCAAAAATAAGGTTATCAAGTGAGAAATCACCATGAGTGACGA<br>CTGAATCCGGTGAGAATGGCAAAAGTTTATGCATTTCTTTCCAGACTTGTT<br>CAACAGGCCAGCCATTACGCTCGTCATCAAAATCACTCGCATCAACCAAAC<br>CGTTATTCATTCGTGATTGCGCCTGAGCGAGACGAAATACGCGATCGCTGT<br>TAAAAGGACAATTACAAACAGGAATCGAATGCAACCGGCGCAGGAACAC<br>TGCCAGCGCATCAACAATATTTTCACCTGAATCAGGATATTCTTCTAATACC<br>TGGAATGCTGTTTTCCCGGGGATCGCAGTGGTGAGTAACCATGCATCATC<br>AGGAGTACGGATAAAATGCTTGATGGTCGGAAGAGGCATAAATTCCGTCA<br>GCCAGTTTAGTCTGACCATCTCATCTGTAACATCATTGGCAACGCTACCTTT<br>GCCATGTTTCAGAAACAACTCTGGCGCATCGGGCTTCCCATACAATCGATA<br>GATTGTCGCACCTGATTGCCCGACATTATCGCGAGCCCATTTATACCCATA<br>TAAATCAGCATCCATGTTGGAATTTAATCGCGGCCTAGAGCAAGACGTTTC<br>CCGTTGAATATGGCTCATAACACCCCTTGTATTACTGTTTATGTAAGCAGA<br>CAGTTTTATTGTTCATGACCAAAATCCCTTAACGTGAGTTTTCGTTCCACTG<br>AGCGTCAGACCCCGTAGAAAAGATCAAAGGATCTTCTTGAGATCCTTTTTT<br>TCTGCGCGTAATCTGCTGCTTGCAAACAAAAAAACCACCGCTACCAGCGGT<br>GGTTTGTTTGCCGGATCAAGAGCTACCAACTCTTTTTCCGAAGGTAACTGG<br>CTTCAGCAGAGCGCAGATACCAAATACTGTCCTTCTAGTGTAGCCGTAGTT<br>AGGCCACCACTTCAAGAACTCTGTAGCACCGCCTACATACCTCGCTCTGCT<br>AATCCTGTTACCAGTGGCTGCTGCCAGTGGCGATAAGTCGTGTCTTACCGG<br>GTTGGACTCAAGACGATAGTTACCGGATAAGGCGCAGCGGTCGGGCTGA<br>ACGGGGGGTTCGTGCACACAGCCCAGCTTGGAGCGAACGACCTACACCG<br>AACTGAGATACCTACAGCGTGAGCTATGAGAAAGCGCCACGCTTCCCGAA<br>GGGAGAAAGGCGGACAGGTATCCGGTAAGCGGCAGGGTCGGAACAGGA<br>GAGCGCACGAGGGAGCTTCCAGGGGGAAACGCCTGGTATCTTTATAGTCC<br>TGTCGGGTTTCGCCACCTCTGACTTGAGCGTCGATTTTTGTGATGCTCGTC<br>AGGGGGGCGGAGCCTATGGAAAAACGCCAGCAACGCGGCCTTTTTACGG<br>TTCCTGGCCTTTTGCTGGCCTTTTGCTCACATGTTCTTTCCTGCGTTATCCCC<br>TGATTCTGTGGATAACCGTATTACCGCCTTTGAGTGAGCTGATACCGCTCG<br>CCGCAGCCGAACGACCGAGCGCAGCGAGTCAGTGAGCGAGGAAGCGGA<br>AGAGCGCCTGATGCGGTATTTTCTCCTTACGCATCTGTGCGGTATTTCACA |

| | |
|---|---|
| | CCGCAATGGTGCACTCTCAGTACAATCTGCTCTGATGCCGCATAGTTAAGC<br>CAGTATACACTCCGCTATCGCTACGTGACTGGGTCATGGCTGCGCCCCGAC<br>ACCCGCCAACACCCGCTGACGCGCCCTGACGGGCTTGTCTGCTCCCGGCAT<br>CCGCTTACAGACAAGCTGTGACCGTCTCCGGGAGCTGCATGTGTCAGAGG<br>TTTTCACCGTCATCACCGAAACGCGCGAGGCAGCTGCGGTAAAGCTCATC<br>AGCGTGGTCGTGAAGCGATTCACAGATGTCTGCCTGTTCATCCGCGTCCA<br>GCTCGTTGAGTTTCTCCAGAAGCGTTAATGTCTGGCTTCTGATAAAGCGGG<br>CCATGTTAAGGGCGGTTTTTTCCTGTTTGGTCACTGATGCCTCCGTGTAAG<br>GGGGATTTCTGTTCATGGGGGTAATGATACCGATGAAACGAGAGAGGAT<br>GCTCACGATACGGGTTACTGATGATGAACATGCCCGGTTACTGGAACGTT<br>GTGAGGGTAAACAACTGGCGGTATGGATGCGGCGGGACCAGAGAAAAAT<br>CACTCAGGGTCAATGCCAGCGCTTCGTTAATACAGATGTAGGTGTTCCACA<br>GGGTAGCCAGCAGCATCCTGCGATGCAGATCCGGAACATAATGGTGCAG<br>GGCGCTGACTTCCGCGTTTCCAGACTTTACGAAACACGGAAACCGAAGAC<br>CATTCATGTTGTTGCTCAGGTCGCAGACGTTTTGCAGCAGCAGTCGCTTCA<br>CGTTCGCTCGCGTATCGGTGATTCATTCTGCTAACCAGTAAGGCAACCCCG<br>CCAGCCTAGCCGGGTCCTCAACGACAGGAGCACGATCATGCGCACCCGTG<br>GGGCCGCCATGCCGGCGATAATGGCCTGCTTCTCGCCGAAACGTTTGGTG<br>GCGGGACCAGTGACGAAGGCTTGAGCGAGGGCGTGCAAGATTCCGAATA<br>CCGCAAGCGACAGGCCGATCATCGTCGCGCTCCAGCGAAAGCGGTCCTCG<br>CCGAAAATGACCCAGAGCGCTGCCGGCACCTGTCCTACGAGTTGCATGAT<br>AAAGAAGACAGTCATAAGTGCGGCGACGATAGTCATGCCCCGCGCCCACC<br>GGAAGGAGCTGACTGGGTTGAAGGCTCTCAAGGGCATCGGTCGAGATCC<br>CGGTGCCTAATGAGTGAGCTAACTTACATTAATTGCGTTGCGCTCACTGCC<br>CGCTTTCCAGTCGGGAAACCTGTCGTGCCAGCTGCATTAATGAATCGGCCA<br>ACGCGCGGGGAGAGGCGGTTTGCGTATTGGGCGCCAGGGTGGTTTTTCTT<br>TTCACCAGTGAGACGGGCAACAGCTGATTGCCCTTCACCGCCTGGCCCTG<br>AGAGAGTTGCAGCAAGCGGTCCACGCTGGTTTGCCCCAGCAGGCGAAAAT<br>CCTGTTTGATGGTGGTTAACGGCGGGATATAACATGAGCTGTCTTCGGTAT<br>CGTCGTATCCCACTACCGAGATATCCGCACCAACGCGCAGCCCGGACTCG<br>GTAATGGCGCGCATTGCGCCCAGCGCCATCTGATCGTTGGCAACCAGCAT<br>CGCAGTGGGAACGATGCCCTCATTCAGCATTTGCATGGTTTGTTGAAAACC<br>GGACATGGCACTCCAGTCGCCTTCCCGTTCCGCTATCGGCTGAATTTGATT<br>GCGAGTGAGATATTTATGCCAGCCAGCCAGACGCAGACGCGCCGAGACA<br>GAACTTAATGGGCCCGCTAACAGCGCGATTTGCTGGTGACCCAATGCGAC<br>CAGATGCTCCACGCCCAGTCGCGTACCGTCTTCATGGGAGAAAATAATACT<br>GTTGATGGGTGTCTGGTCAGAGACATCAAGAAATAACGCCGGAACATTAG<br>TGCAGGCAGCTTCCACAGCAATGGCATCCTGGTCATCCAGCGGATAGTTA<br>ATGATCAGCCCACTGACGCGTTGCGCGAGAAGATTGTGCACCGCCGCTTT<br>ACAGGCTTCGACGCCGCTTCGTTCTACCATCGACACCACCACGCTGGCACC<br>CAGTTGATCGGCGCGAGATTTAATCGCCGCGACAATTTGCGACGGCGCGT<br>GCAGGGCCAGACTGGAGGTGGCAACGCCAATCAGCAACGACTGTTTGCCC<br>GCCAGTTGTTGTGCCACGCGGTTGGGAATGTAATTCAGCTCCGCCATCGCC<br>GCTTCCACTTTTTCCCGCGTTTTCGCAGAAACGTGGCTGGCCTGGTTCACC<br>ACGCGGGAAACGGTCTGATAAGAGACACCGGCATACTCTGCGACATCGTA<br>TAACGTTACTGGTTTCACATTCACCACCCTGAATTGACTCTCTTCCGGGCGC<br>TATCATGCCATACCGCGAAAGGTTTTGCGCCATTCGATGGTGTCCGGGATC<br>TCGACGCTCTCCCTTATGCGACTCCTGCATTAGGAAGCAGCCCAGTAGTAG |

| | |
|---|---|
| | GTTGAGGCCGTTGAGCACCGCCGCCGCAAGGAATGGTGCATGCAAGGAG<br>ATGGCGCCCAACAGTCCCCCGGCCACGGGGCCTGCCACCATACCCACGCC<br>GAAACAAGCGCTCATGAGCCCGAAGTGGCGAGCCCGATCTTCCCCATCGG<br>TGATGTCGGCGATATAGGCGCCAGCAACCGCACCTGTGGCGCCGGTGATG<br>CCGGCCACGATGCGTCCGGCGTAGAGGATCGAGATCTCGATCCCGCGAAA<br>TTAATACGACTCACTATAGGGGAATTGTGAGCGGATAACAATTCCCCTCTA<br>GAAATAATTTTGTTTAACTTTAAGAAGGAGATATACCATGGGCAGCAGCC<br>ATCATCATCATCATCACAGCAGCGGCCTGGTGCCGCGCGGCAGCCATATG |
| JRF_A12-13_R | CAGGCCGCTGCTGTG |
| JRF_A12-13_F | CTCGAGCACCACCACCAC |

As for our previous studies, the plasmids will be deposited at Addgene.

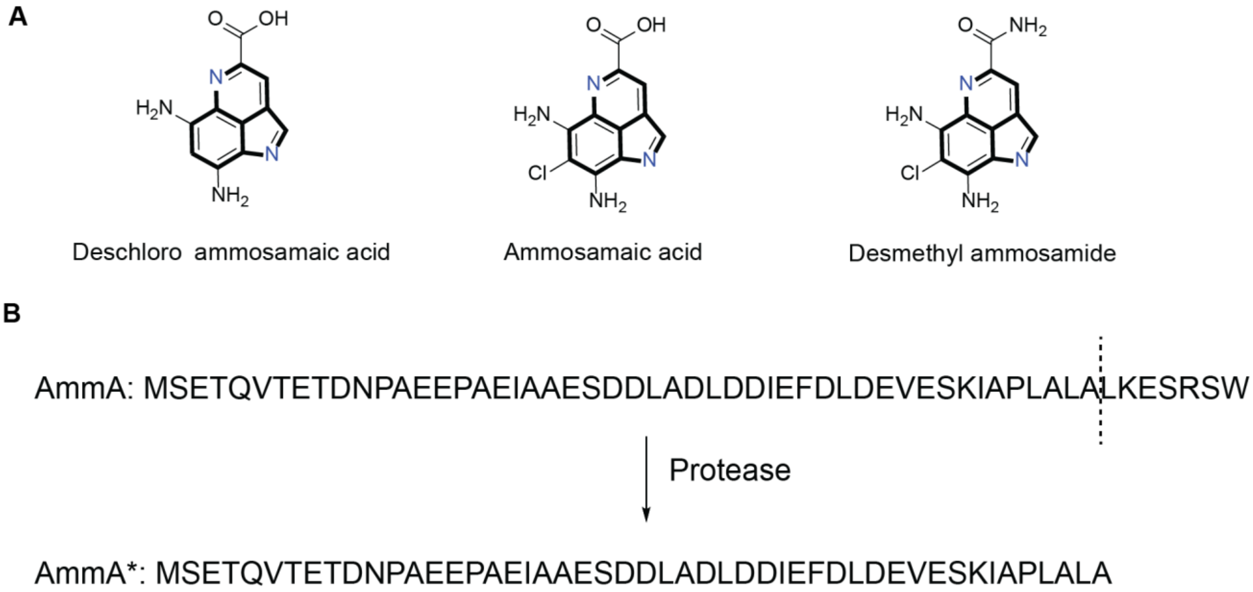


**Figure S1.** (A) Pyrroloiminoquinone metabolites proposed and identifiedfrom ammosamide-related BGCs. Ammosamaic acid was identified from a strain in which the gene for Amm3 (chlorinase) was disrupted and desmethyl ammosamide was identified from a strain in which the gene for Amm23 (methyltransferase) was disrupted. Deschloro ammosamaic acid was proposed as an intermediate in the lymphostin BGC. (B) Precursor peptide sequence AmmA and AmmA* obtained after protease processing.[5]

**Table S2.** ESI-MS/MS analysis of in vivo produced intermediate **7** digested with trypsin, corresponding to Figure 2C. Experimental and theoretical values represent $[M+H]^+$ ions.

| Fragment Type | Fragmented Bond Number | Neutral Loss | Experimental m/z | Theoretical m/z | Mass Error (ppm) |
|---|---|---|---|---|---|
| a | 1 | | 86.1 | 86.1 | 5.6 |
| a | 2 | | 157.1 | 157.1 | 6.1 |
| y | 1 | -NH3 | 158.1 | 158.1 | 8.2 |
| y | 1 | | 175.1 | 175.1 | 6.6 |
| b | 2 | | 185.1 | 185.1 | 3.5 |
| b | 3 | | 282.2 | 282.2 | 3.1 |
| a | 4 | | 367.3 | 367.3 | 3.4 |
| y | 2 | -NH3 | 388.2 | 388.2 | 3.8 |
| b | 4 | | 395.3 | 395.3 | 4.4 |
| y | 2 | | 405.2 | 405.2 | 2.8 |
| a | 5 | | 438.3 | 438.3 | 1.0 |
| y | 3 | -NH3 | 459.2 | 459.2 | 1.2 |
| b | 5 | | 466.3 | 466.3 | 2.0 |
| y | 3 | | 476.2 | 476.2 | 1.9 |
| a | 6 | | 551.4 | 551.4 | -1.5 |
| y | 4 | -NH3 | 572.3 | 572.3 | 1.9 |
| b | 6 | | 579.4 | 579.4 | 0.8 |
| y | 4 | | 589.3 | 589.3 | -0.2 |
| y | 5 | -NH3 | 643.3 | 643.3 | 5.6 |
| b | 7 | | 650.4 | 650.4 | 2.8 |
| y | 5 | | 660.4 | 660.4 | -3.0 |
| a | 8 | | 852.5 | 852.5 | 1.3 |
| y | 7 | -NH3 | 853.5 | 853.5 | 7.2 |
| y | 7 | | 870.5 | 870.5 | -3.2 |
| b | 8 | | 880.5 | 880.5 | 3.0 |
| M+H | | -NH3 | 1037.6 | 1037.6 | 2.9 |
| M+H | | | 1054.6 | 1054.6 | 2.2 |

**A**

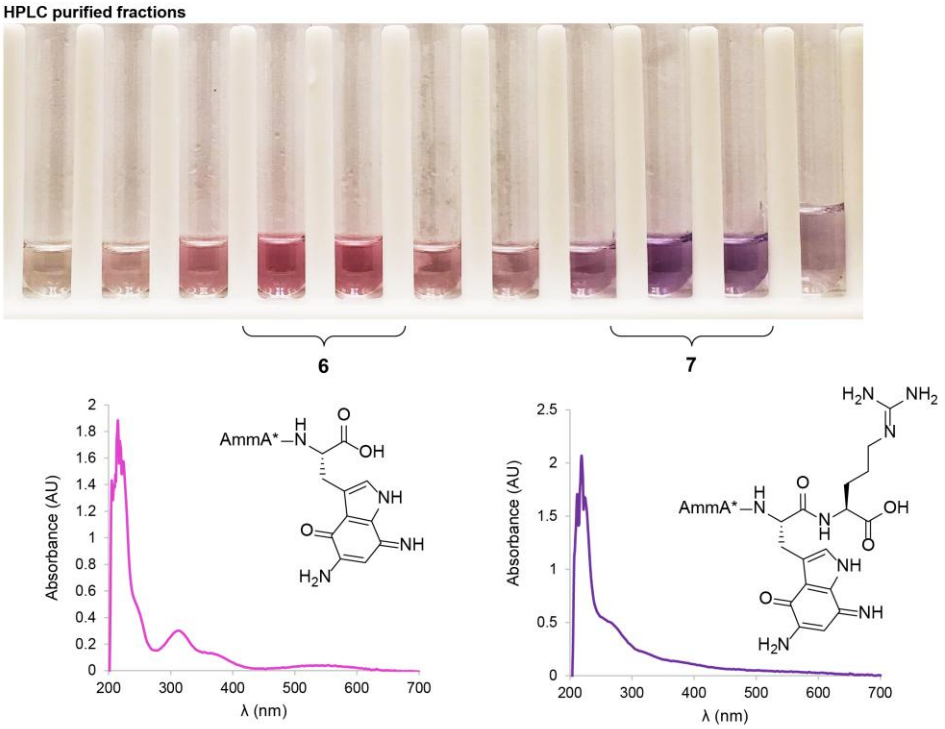

HPLC purified fractions
6
7
Absorbance (AU)
λ (nm)
AmmA*
NH
OH
O
H₂N
NH₂


**B**

**TOCSY**

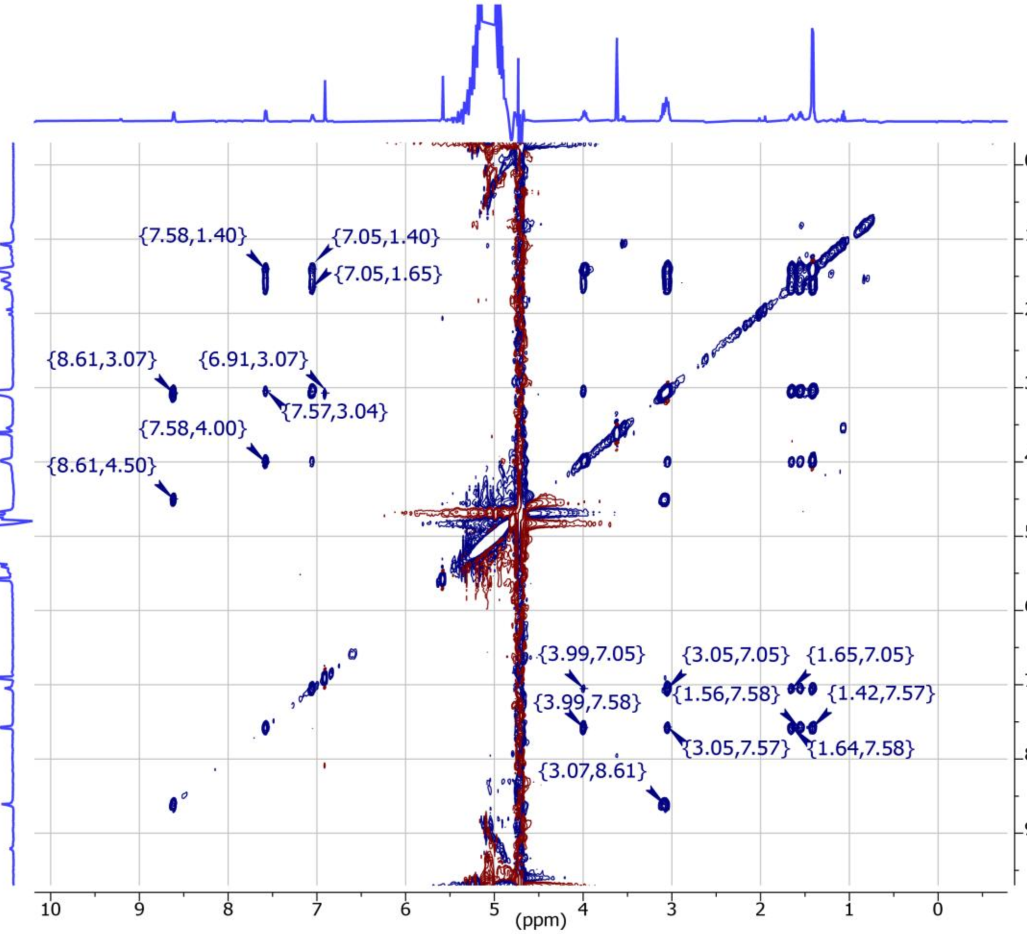

{7.58,1.40}
{7.05,1.40}
{7.05,1.65}
{8.61,3.07}
{6.91,3.07}
{7.57,3.04}
{7.58,4.00}
{8.61,4.50}
{3.99,7.05}
{3.05,7.05}
{1.65,7.05}
{1.56,7.58}
{1.42,7.57}
{3.99,7.58}
{3.05,7.57}
{1.64,7.58}
{3.07,8.61}
(ppm)

C

NOESY

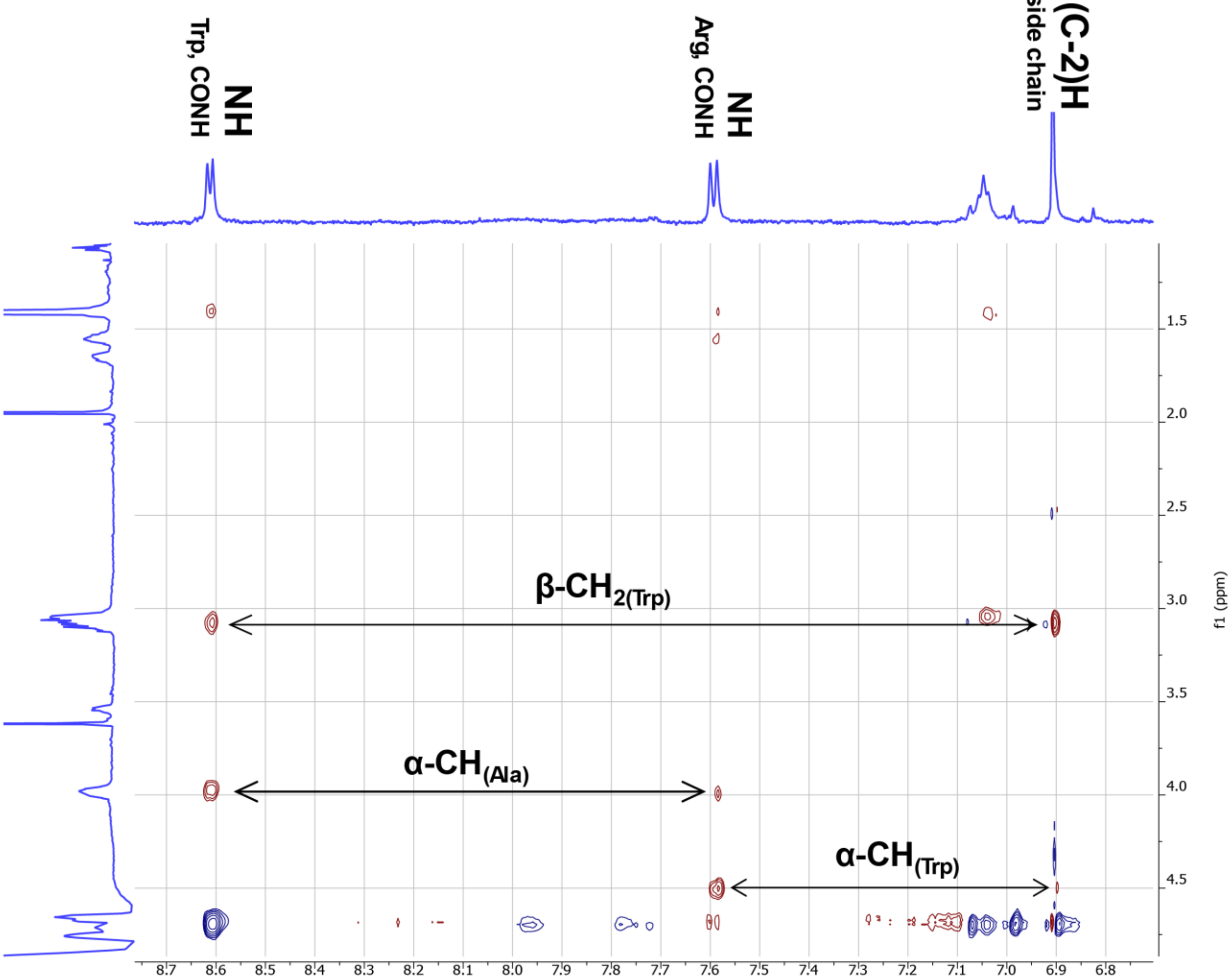


D

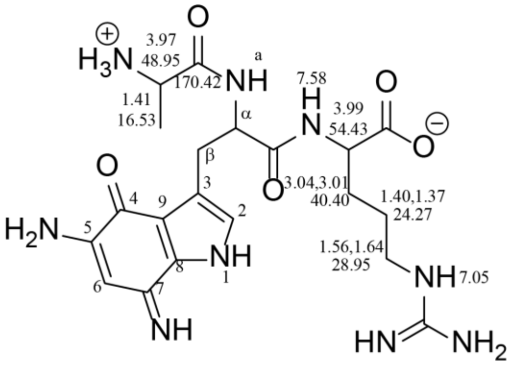


E

| Trp | $\delta_H$ (ppm) | $\delta_C$ (ppm) |
|---|---|---|
| a | 8.61 | n.a. |
| α | 4.5 | n.o. |
| β | 3.07 | 27.17 |
| 2 | 6.91 | 126.5 |
| 3 | n.a. | 121.92 |
| 4 | n.a. | 177.32 |
| 5 | n.o. | n.o. |
| 6 | 5.57 | 89.94 |
| 7 | n.o. | n.o. |
| 8 | n.a. | 119.62 |
| 9 | n.a. | 128.65 |

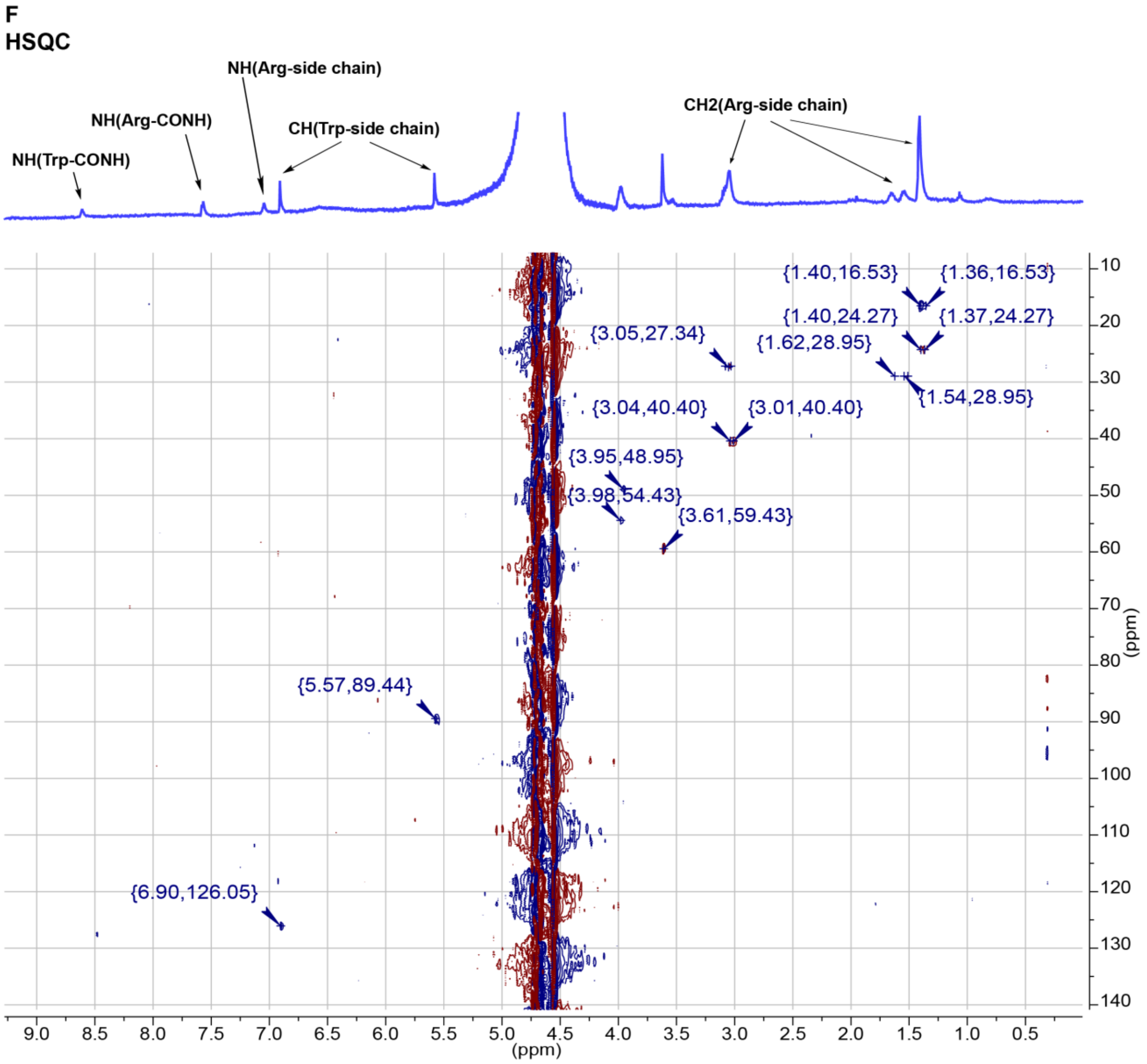


**Figure S2.** Purification and characterization of compound **7**. (A) Illustration of the change in color upon adding Arg to peptide **6**. (B) 2D NMR analysis (TOCSY) of a chymotrypsin fragment of intermediate **7** in 10% $D_2O$, using a $D_2O$ matched Shigemi tube. (C) 2D NMR analysis (NOESY) of a chymotrypsin fragment of intermediate **7** in 10% $D_2O$, using a $D_2O$ matched Shigemi tube. (D) Chemical shifts of the chymotrypsin fragment of **7**. (E) Observed chemical shifts and assignments for the modified Trp. n.a.=not applicable, n.o.=not observed. (F) Heteronuclear Single Quantum Coherence (HSQC) spectrum of a chymotrypsin fragment of intermediate **7** (panel D) in 10% $D_2O$, using a $D_2O$ matched Shigemi tube.

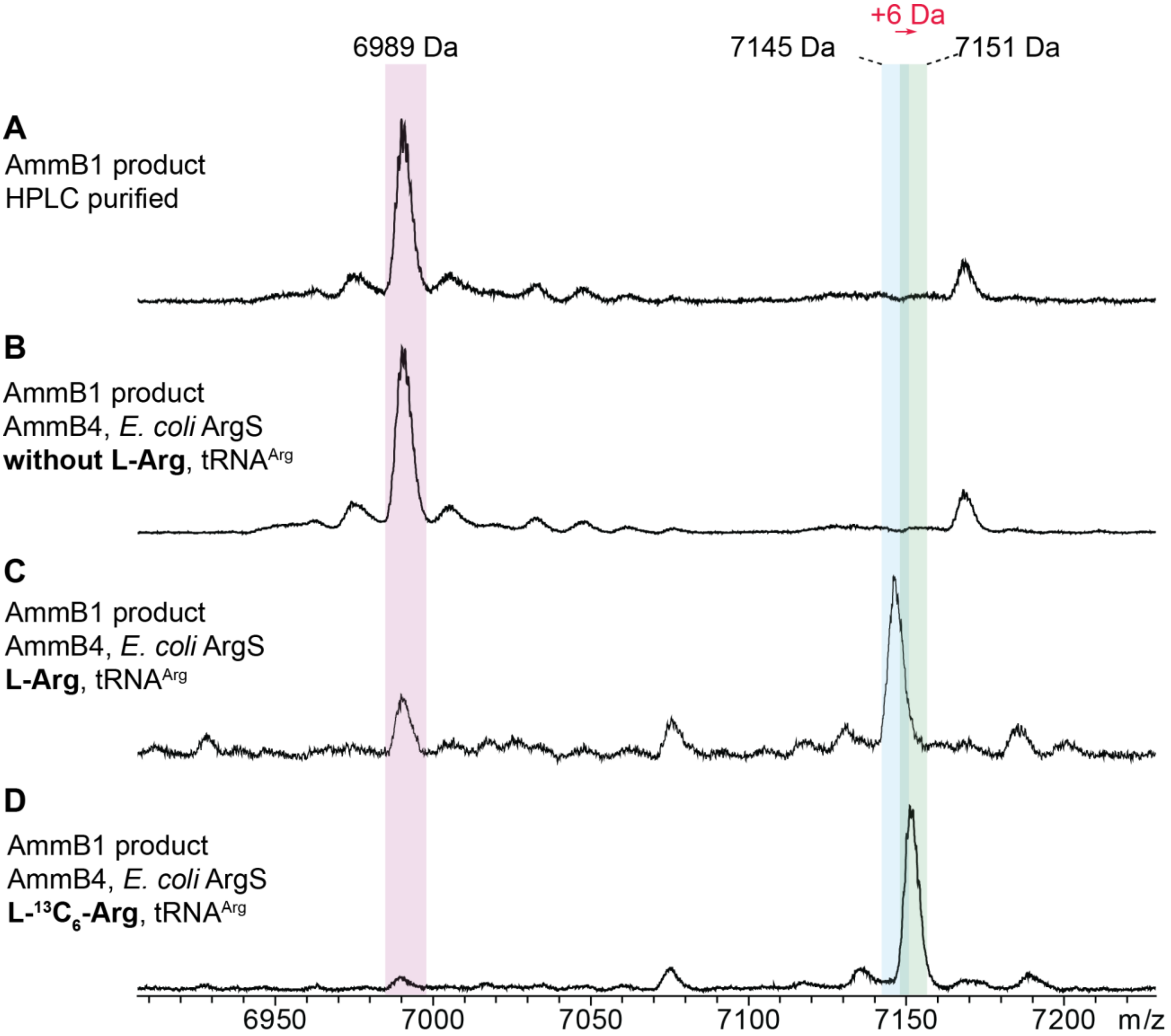


**Figure S3.** AmmB$_4$ reaction using purified tRNA$^{Arg}$, *E. coli* ArgRS, HPLC purified substrate (intermediate **6**) with and without L-Arg. (A) Intermediate **6** reference compound as purified by HPLC. Calculated average m/z for $[M+H]^+$ for **6**: 6990 Da, observed average m/z for $[M+H]^+$ for **6**: 6989 Da. (B) In vitro reaction using tRNA$^{Arg}$, *E. coli* ArgRS, AmmB$_4$, intermediate **6** but without L-Arg. (C) In vitro reaction using tRNA$^{Arg}$, *E. coli* ArgRS, AmmB$_4$, intermediate **6**, and L-Arg. Calculated average m/z for $[M+H]^+$ for **7**: 7146 Da, observed average m/z for $[M+H]^+$ for **7**: 7145 Da. (D) In vitro reaction using tRNA$^{Arg}$, *E. coli* ArgRS, AmmB$_4$, intermediate **6**, and L-$^{13}C_6$-Arg. Calculated average m/z for $[M+H]^+$ for **7**: 7152 Da, observed average m/z for $[M+H]^+$ for **7**: 7151 Da.

A

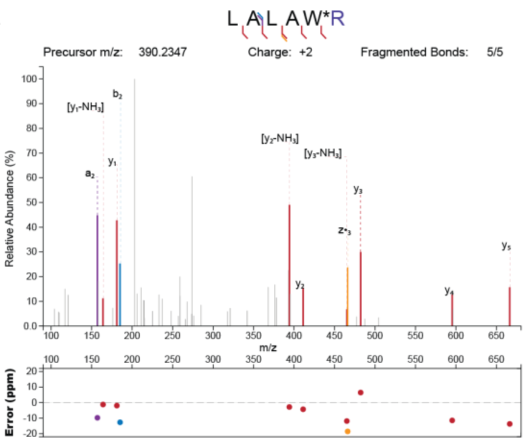


| Fragment Type | Fragmented Bond Number | Neutral Loss | Experimental Mz | Theoretical Mz | Mass Error (ppm) |
|---|---|---|---|---|---|
| a | 2 | | 157.132 | 157.1335 | -9.7742 |
| y | 1 | -NH3 | 164.1126 | 164.1122 | 2.4249 |
| y | 1 | | 181.139 | 181.1388 | 1.3738 |
| b | 2 | | 185.1261 | 185.1285 | -12.6964 |
| y | 2 | | 206.1129 | 206.1132 | -1.3042 |
| y | 2 | -NH3 | 394.1921 | 394.1925 | -1.0453 |
| y | 2 | | 411.218 | 411.2191 | -2.5805 |
| y | 3 | -NH3 | 465.2248 | 465.2296 | -10.3735 |
| y | 3 | | 482.26 | 482.2562 | 7.9312 |
| y | 4 | | 595.3341 | 595.3402 | -10.3053 |
| y | 5 | | 666.3689 | 666.3773 | -12.6792 |

B

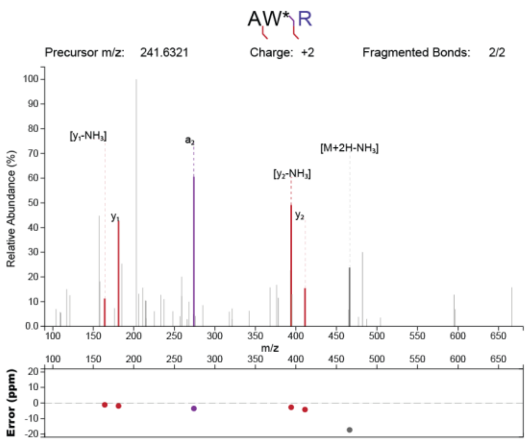


| Fragment Type | Fragmented Bond Number | Neutral Loss | Experimental Mz | Theoretical Mz | Mass Error (ppm) |
|---|---|---|---|---|---|
| y | 1 | -NH3 | 164.1126 | 164.1122 | 2.4249 |
| y | 1 | | 181.139 | 181.1388 | 1.3738 |
| y | 2 | | 206.1129 | 206.1132 | -1.3042 |
| a | 2 | | 274.1289 | 274.1298 | -3.2315 |
| y | 2 | -NH3 | 394.1921 | 394.1925 | -1.0453 |
| y | 2 | | 411.218 | 411.2191 | -2.5805 |
| M+2H | | -NH3 | 466.2295 | 466.2369 | -15.8772 |

C

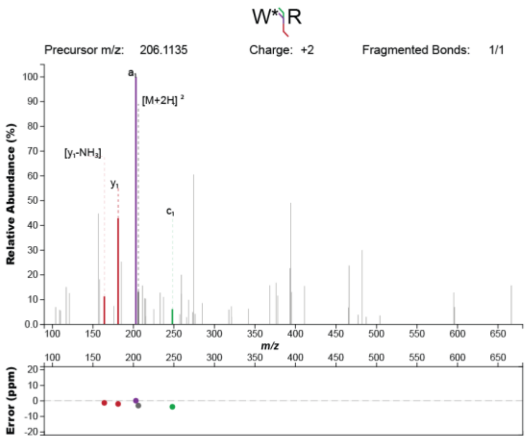


| Fragment Type | Fragmented Bond Number | Neutral Loss | Experimental Mz | Theoretical Mz | Mass Error (ppm) |
|---|---|---|---|---|---|
| y | 1 | -NH3 | 164.1126 | 164.1122 | 2.4249 |
| y | 1 | | 181.139 | 181.1388 | 1.3738 |
| a | 1 | | 203.0928 | 203.0927 | 0.631 |
| M+2H | | | 206.1129 | 206.1132 | -1.3042 |

**Figure S4.** ESI-MS/MS analysis of the $[M+2H]^{2+}$ ion for the C-terminal modified intermediate **8** with labeled L-$^{13}C_6$-Arg and digested with proalanase. (A) Fragmentation pattern for LALAW*R, where R is $^{13}C_6$-Arg. (B) Additional fragmentation of 390 Da ion produces AW*R, where R is also $^{13}C_6$-isotopically labeled. (C) Additional fragmentation of 206 Da ion produces W*R, where R is also $^{13}C_6$-isotopically labeled.

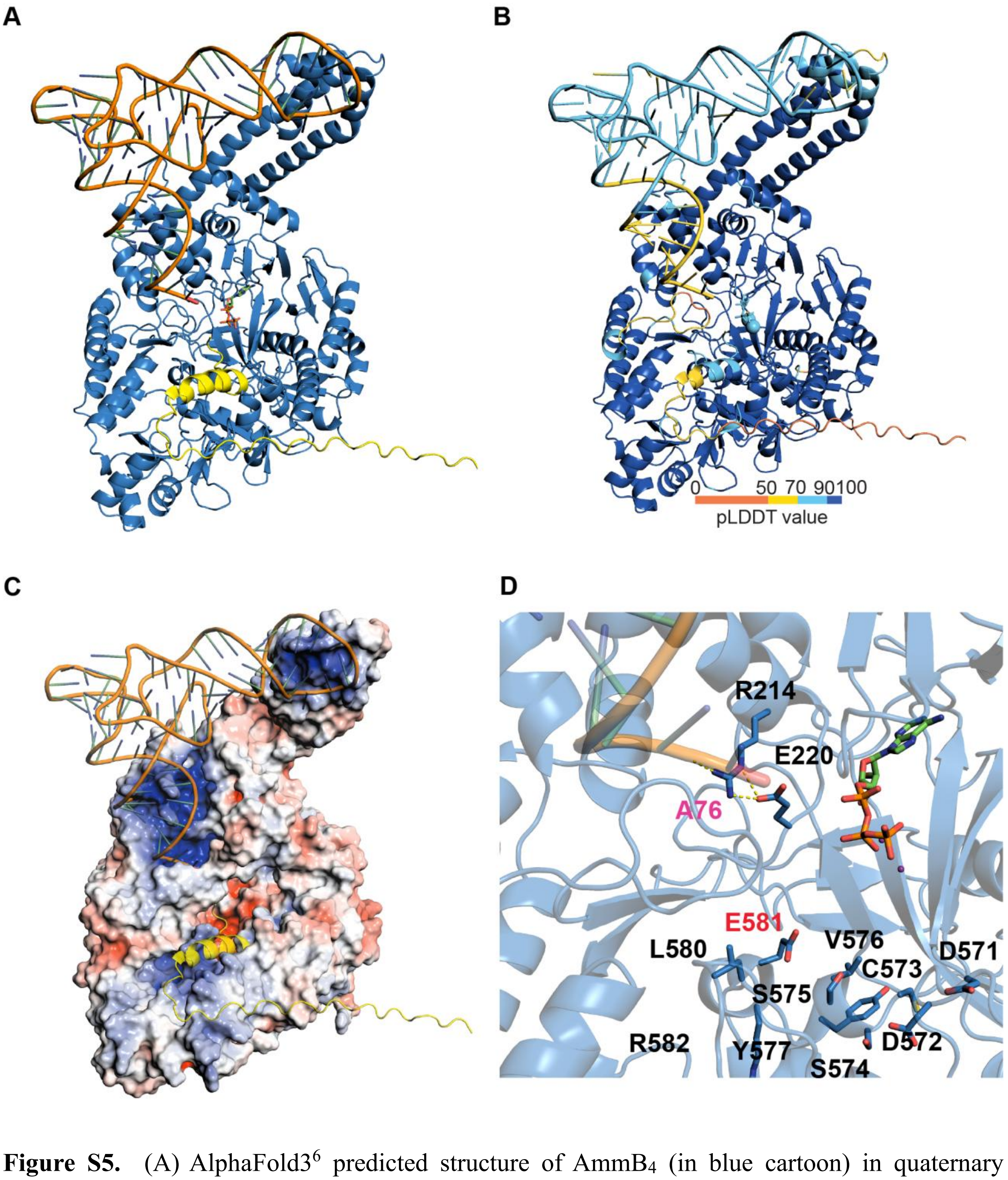


**Figure S5.** (A) AlphaFold3[6] predicted structure of AmmB$_4$ (in blue cartoon) in quaternary complex with tRNA$^{Arg}$ (in orange ribbon), AmmA*W (in yellow cartoon), ATP (in green sticks), and two $Mg^{2+}$ ions (in purple spheres) visualized by PyMOL. (B) AlphaFold3 predicted structure of AmmB$_4$ in quaternary complex colored by predicted local distance difference test (pLDDT) scores. (C) Electrostatic surface potential calculated for AmmB$_4$. Positive potential (blue color) overlaps with tRNA contacts as well as some parts of the peptide substrate (in yellow). Negative potential (red color) is observed in the enzyme active site near the C-terminus of the peptide substrate (yellow) and the A76 nucleotide that forms the 3' terminus of $tRNA^{Arg}$. (D) Zoom-in showing Glu581 located in a helix positioned between the CCA and ATP binding site, Glu581 can tentatively interact with the side chain of the Arg attached to the tRNA. Glu220 is conserved in all PEARLs and is unlikely to provide recognition of the Arg side chain.

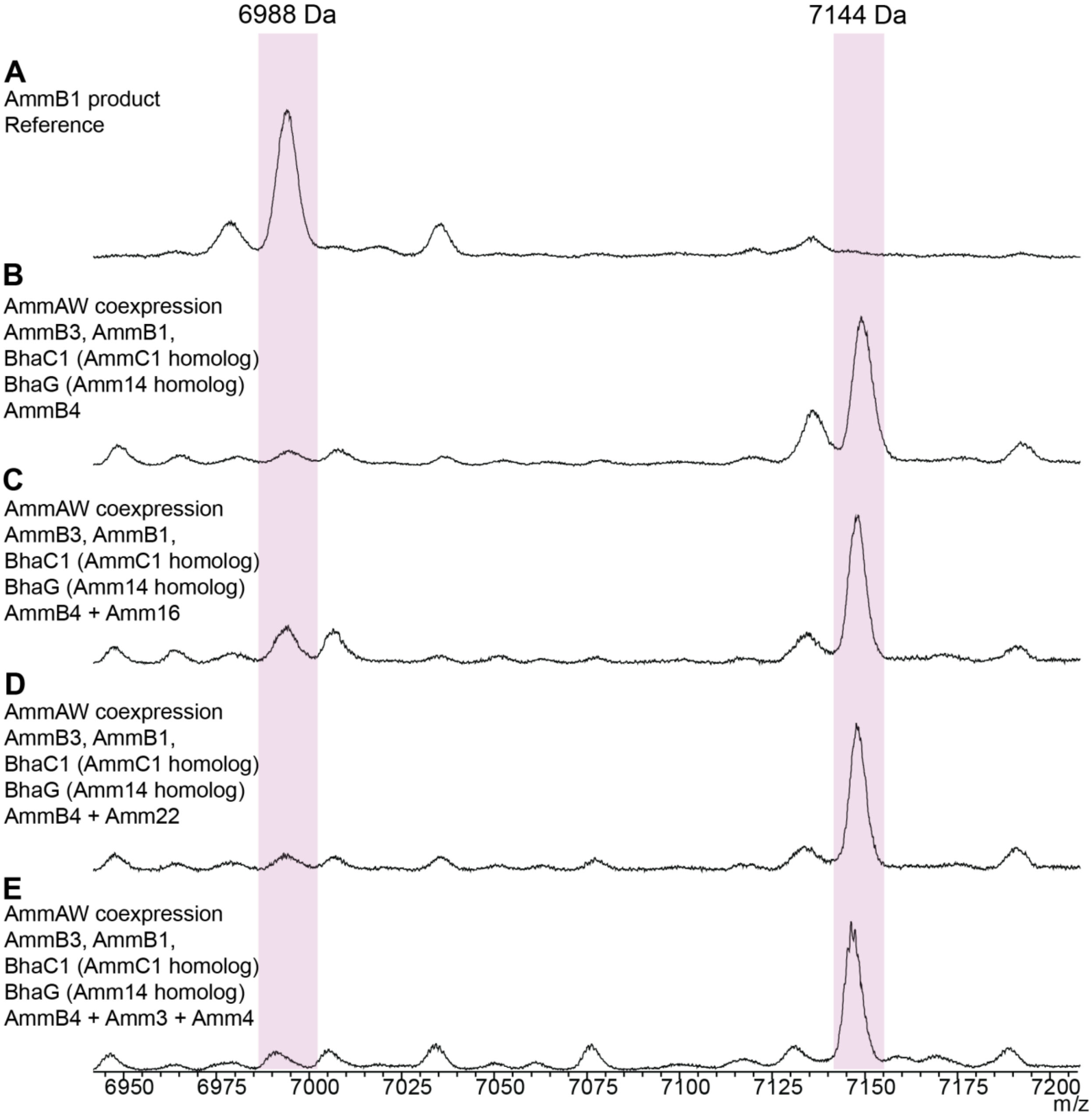


**Figure S6.** MALDI-TOF mass spectra of co-expressions with various enzymes (indicated in red font below) that did not lead to additional mass changes after modification by $AmmB_4$. (A) Peptide obtained after HPLC purification of the product from co-expression of AmmA*W, $AmmB_3$, $BhaC_1$ (AmmC1 homolog), BhaG (Amm14 homolog), and $AmmB_1$. (B) Product of co-expression of

AmmA*W, $AmmB_3$, $BhaC_1$ ($AmmC_1$ homolog), BhaG (Amm14 homolog), $AmmB_1$, and $AmmB_4$. (C) Product of co-expression of AmmA*W, $AmmB_3$, $BhaC_1$ ($AmmC_1$ homolog), BhaG (Amm14 homolog), $AmmB_1$, $AmmB_4$, and **Amm16**. (D) Product of co-expression of AmmA*W, $AmmB_3$, $BhaC_1$ ($AmmC_1$ homolog), BhaG (Amm14 homolog), $AmmB_1$, $AmmB_4$, and **Amm22**. (E) Product of co-expression of AmmA*W, $AmmB_3$, $BhaC_1$ ($AmmC_1$ homolog), BhaG (Amm14 homolog), $AmmB_1$, $AmmB_4$, and **Amm3 and Amm4**.

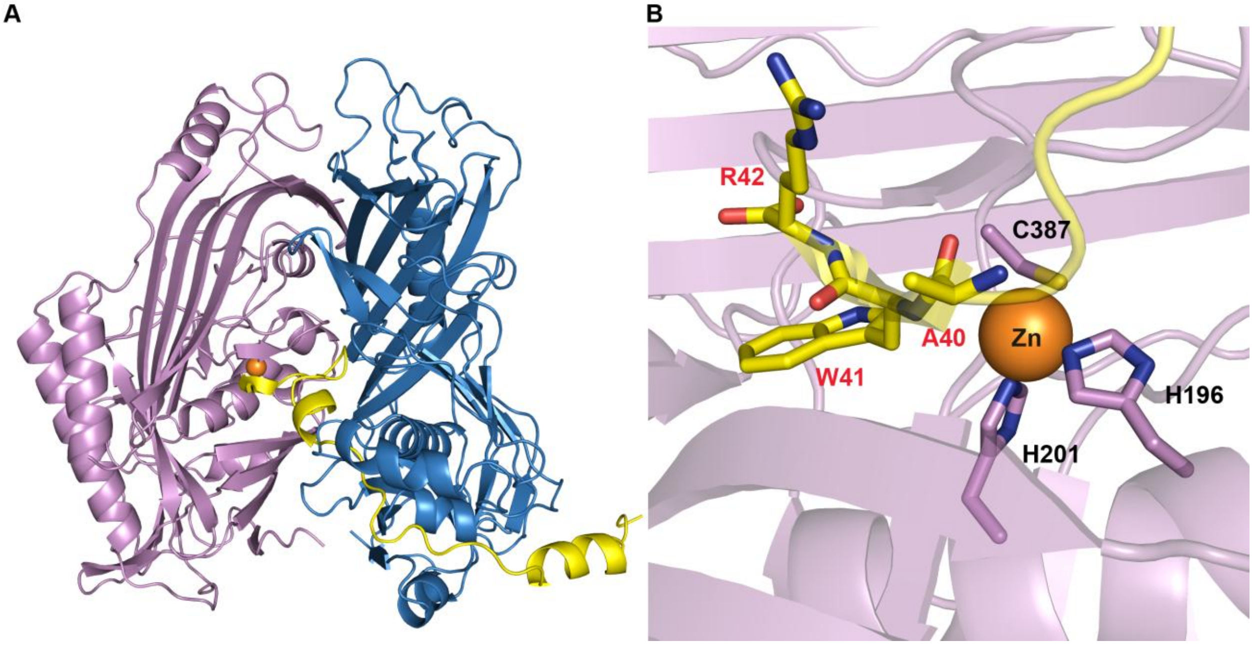


**Figure S7.** AlphaFold3 predicted structure of the Amm12/Amm13 complex with AmmA*WR and $Zn^{2+}$. (A) Quaternary complex of Amm12 (pink) and Amm13 (blue) with a putative interaction with the C-terminus of AmmA*WR (yellow) and Amm12 through a beta sheet interaction proximal to the $Zn^{2+}$ ion (in orange). (B) Predicted $Zn^{2+}$ binding site involves interactions with His196, His201, and Cys387 residues in Amm12. The amide carbonyl bond between Ala40 and Trp41 is observed near the $Zn^{2+}$ ion as expected for C-terminal dipeptide cleavage.

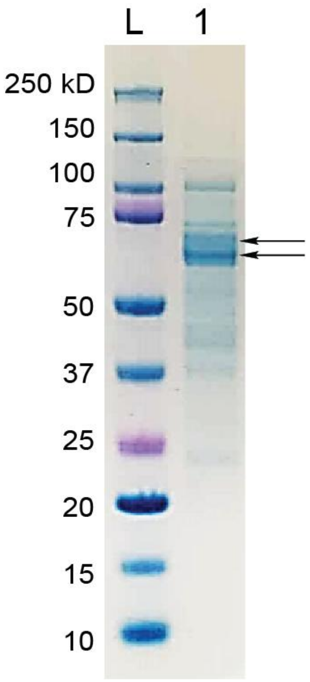


**Figure S8.** SDS-PAGE for the expression of His-tagged-Amm12/Amm13 after size exclusion chromatography.

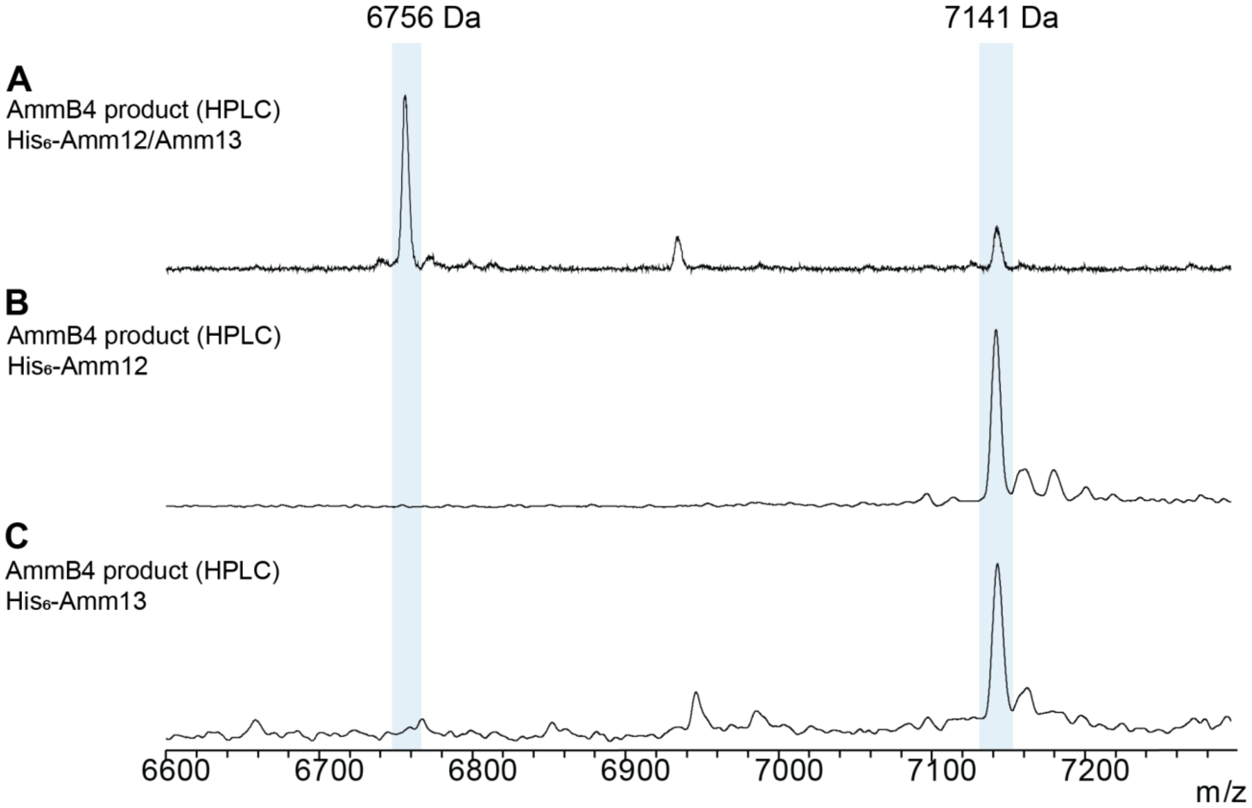


**Figure S9.** MALDI-TOF mass spectra of in vitro reactions of intermediate **7** with (A) His-tagged Amm12 and pulled down Amm13, (B) His-tagged Amm12, (C) and His-tagged Amm13.

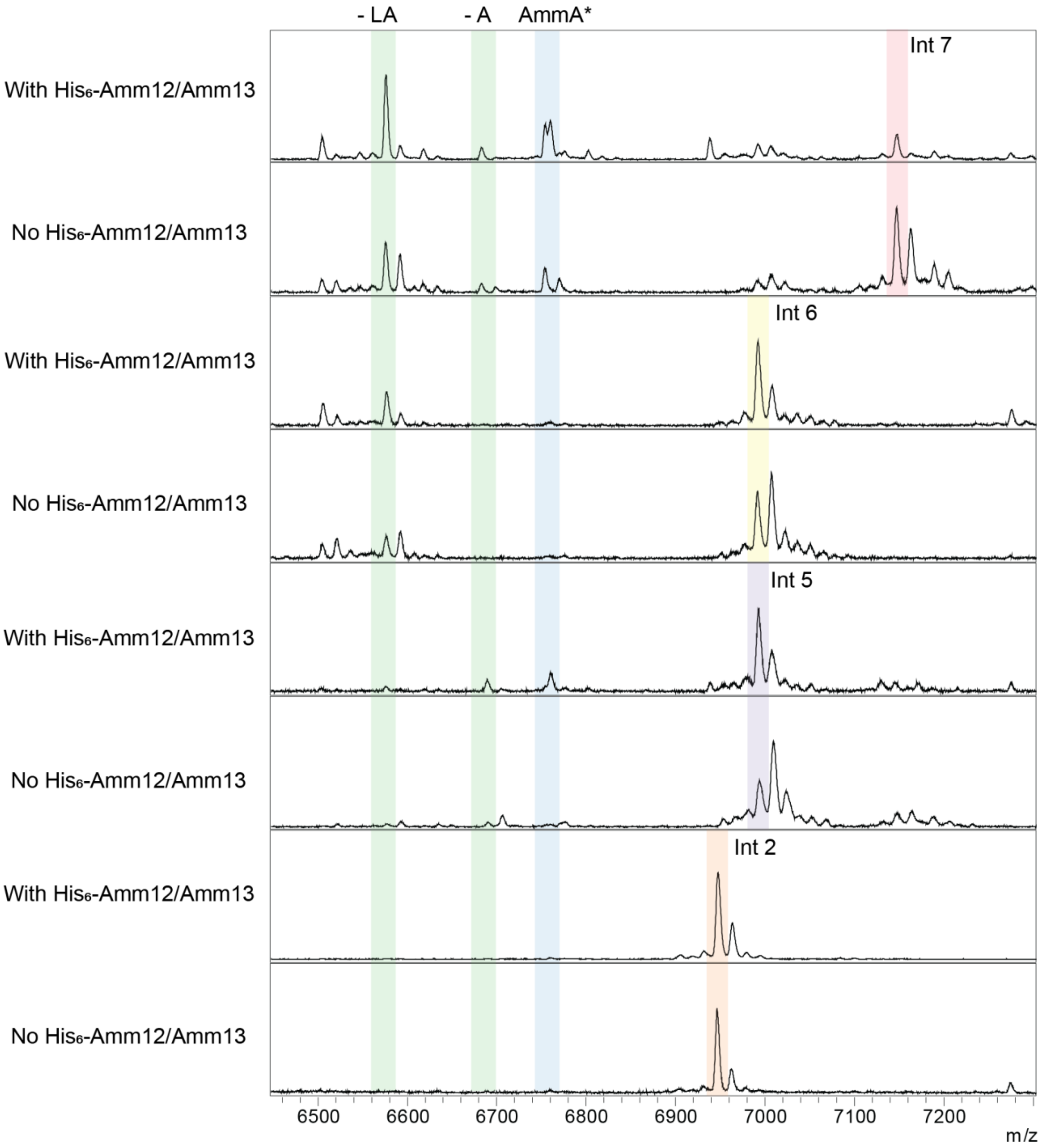


**Figure S10.** Substrate recognition analysis for Amm12/Amm13 activity using intermediates **2**, **5**, **6**, and **7**. While peptide **7** was mostly processed producing AmmA* with corresponding loss of dipeptide **8**, additional removal of the dipeptide Leu-Ala is also observed. All other intermediates that lack the C-terminal Arg residue are not recognized by Amm12/Amm13 for C-terminal proteolysis.

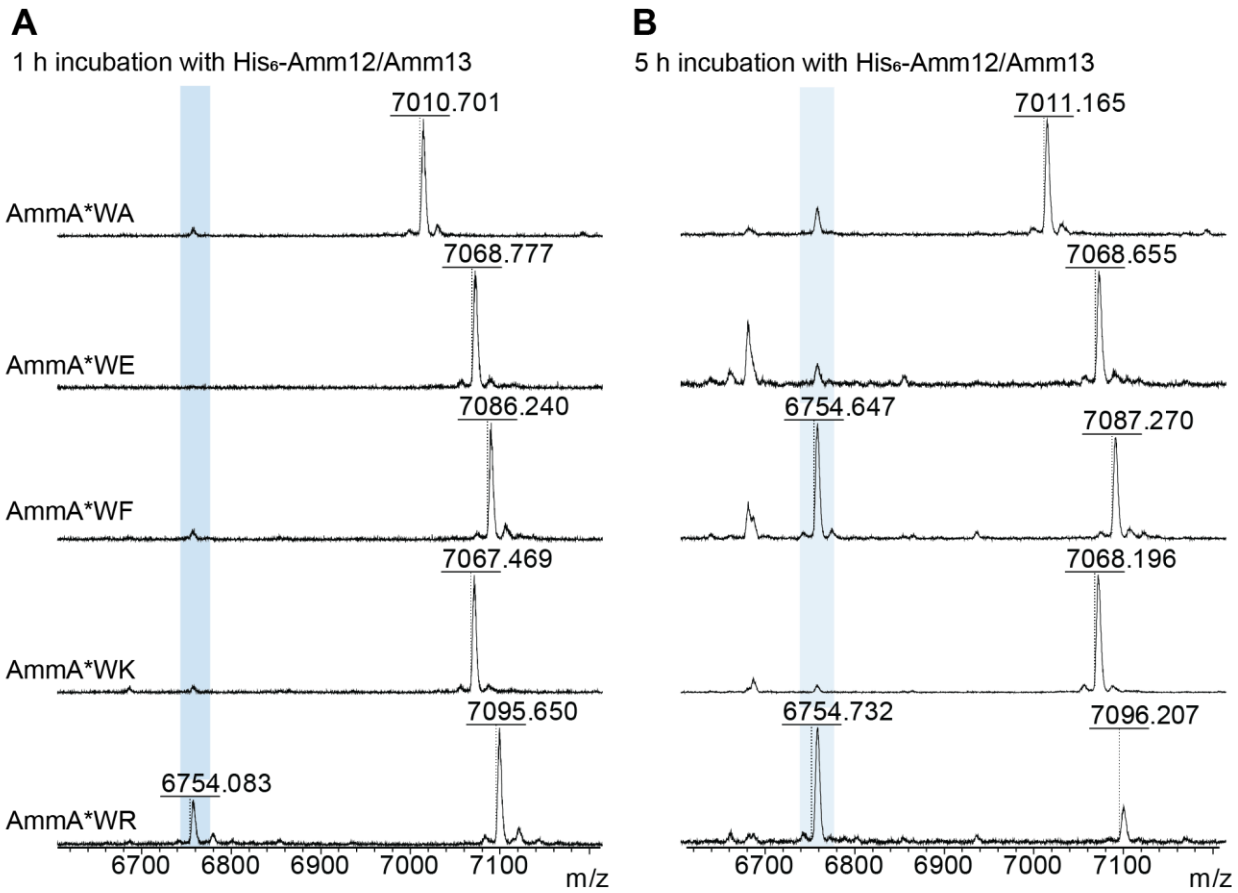


**Figure S11.** Amm12/Amm13 substrate scope. Several analogs of intermediate **7** (AmmA*WA, AmmA*WE, AmmA*WF, AmmA*WK, and AmmAW*R) were expressed and purified as His-tagged fusion constructs. All peptides lacked the posttranslational modifications at the Trp that are preent in **7**. (A) After 1 h incubation at 37 °C with Amm12/Amm13, only analog AmmA*WR was partially cleaved with removal or WR and formation of AmmA*. (B) After 5 h incubation, AmmA*WF was partially cleaved, AmmA*WR was almost fully processed, while AmmA*WA, AmmA*WE, and AmmA*WK were only slightly digested.

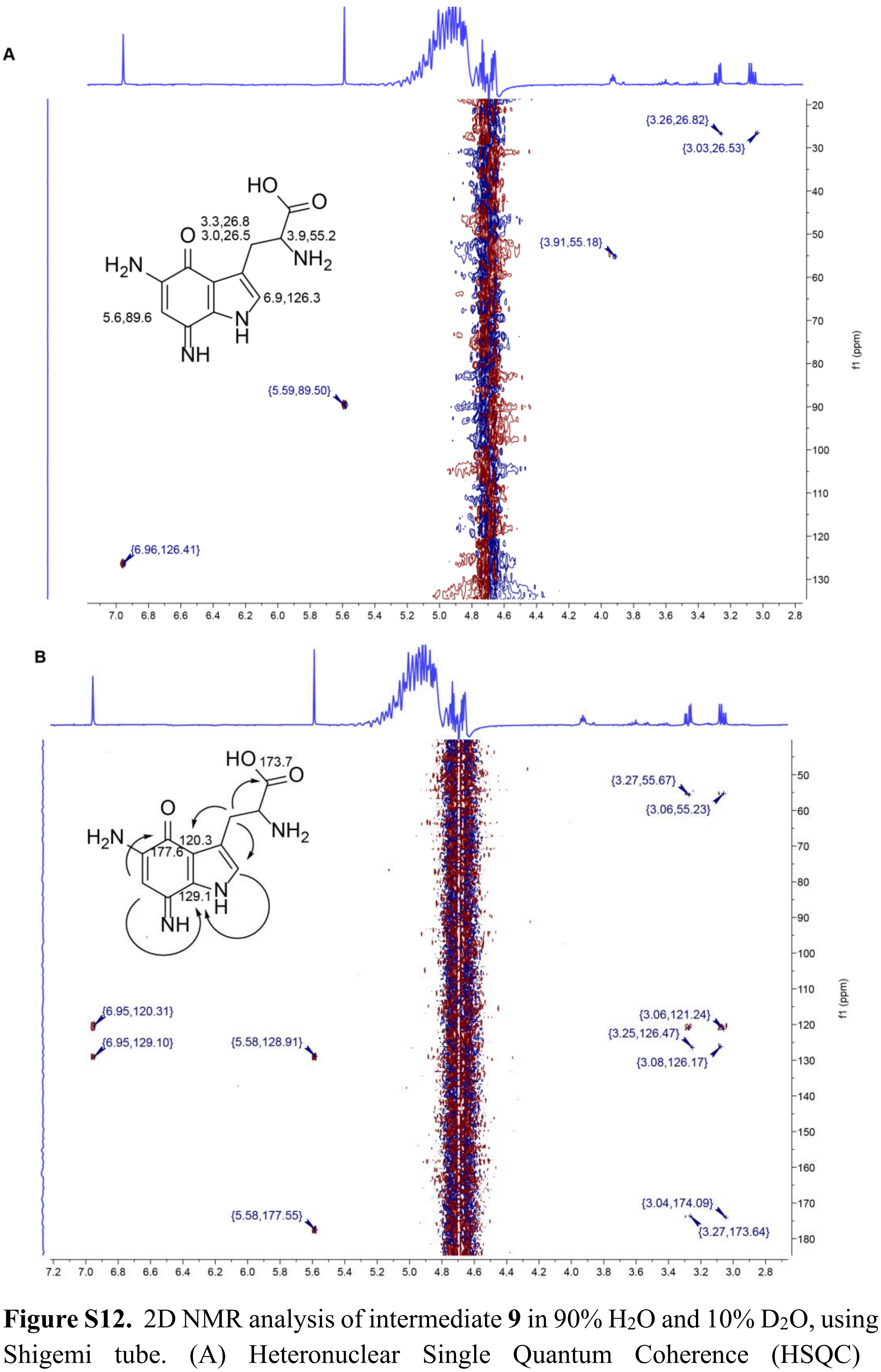


**Figure S12.** 2D NMR analysis of intermediate **9** in 90% $H_2O$ and 10% $D_2O$, using a $D_2O$ matched Shigemi tube. (A) Heteronuclear Single Quantum Coherence (HSQC) spectrum. (B) Heteronuclear Multiple Bond Correlation (HMBC) spectrum. Arrows indicate the correlation between protons and corresponding carbons three-bonds away.

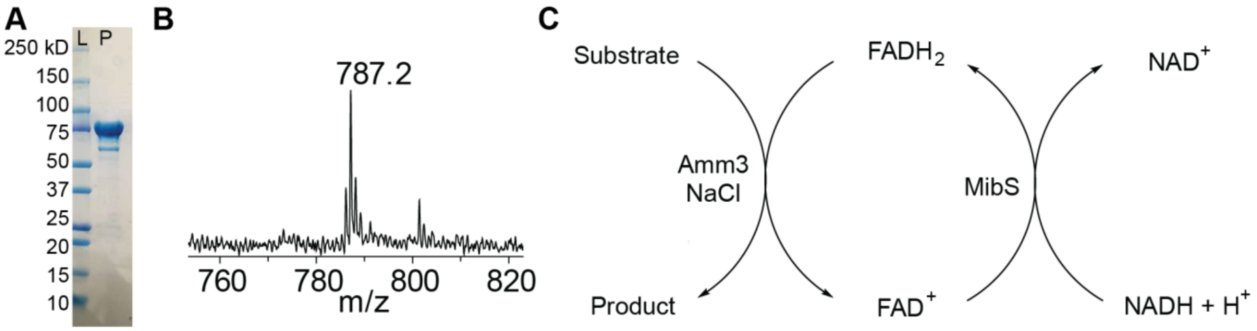


**Figure S13.** (A) SDS-PAGE gel obtained for His-tagged Amm3 coepressed with chaperones as mentioned in the methods description. Molecular weight for His-tagged Amm3 is 74 kDa. (B) MALDI-TOF spectrum of the cofactor released after boiling an aliquot of purified Amm3. Expected mass for FAD: 785.2 Da, observed mass: 787.2 Da. (C) In vitro cascade reaction for Amm3 activity. MibS copurifies with bound FAD which can be reduced using NADH. The reduced flavin, $FADH_2$, $O_2$ and $Cl^-$ form HOCl, the activated form required for Amm3 chlorination of accepted substrates.

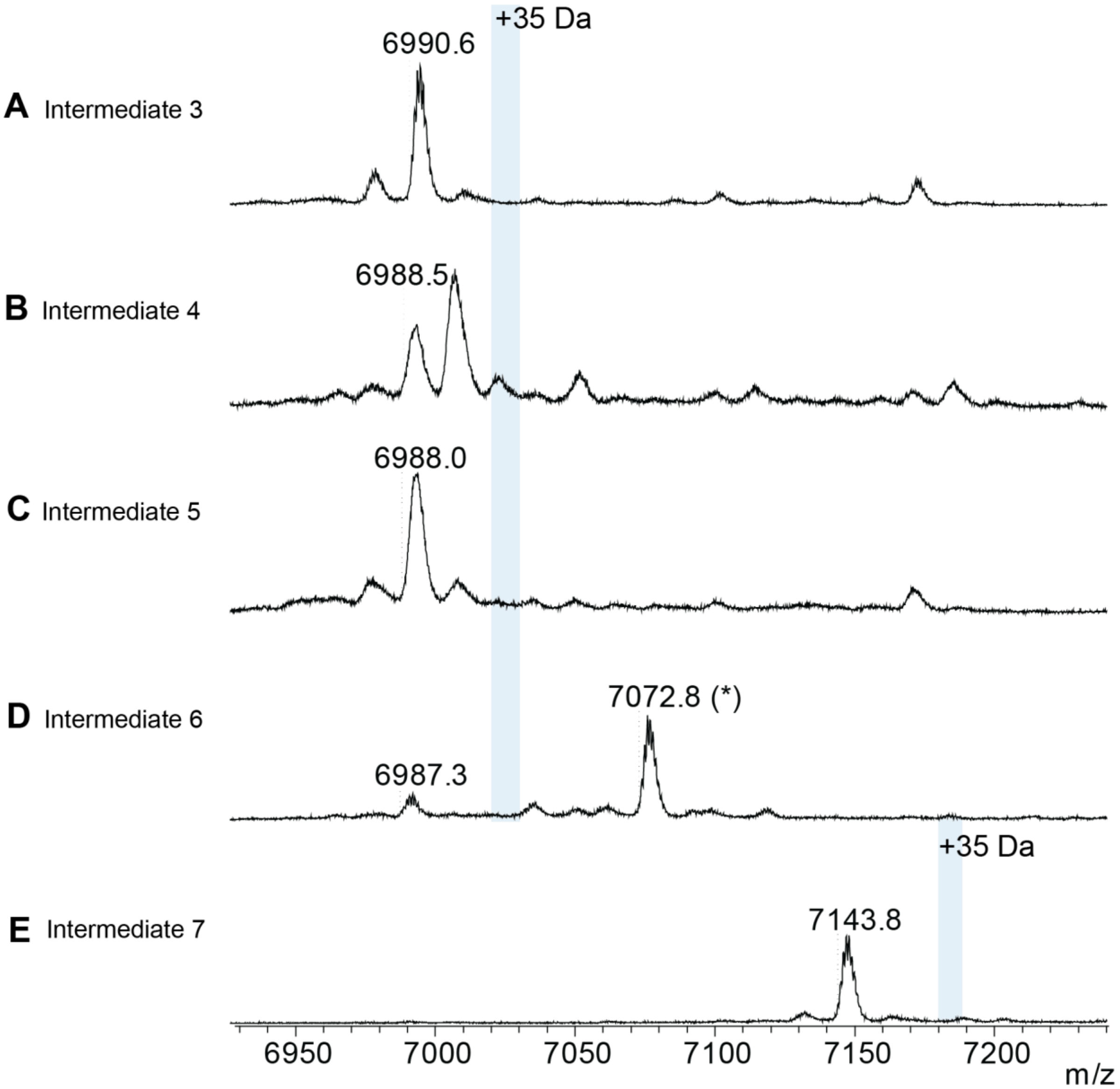


**Figure S14.** MALDI-TOF mass spectra of the products of various peptides incubated with Amm3 in vitro. * indicates diacetylated intermediate **6** produced due to further processing during *E. coli* expression.[1]

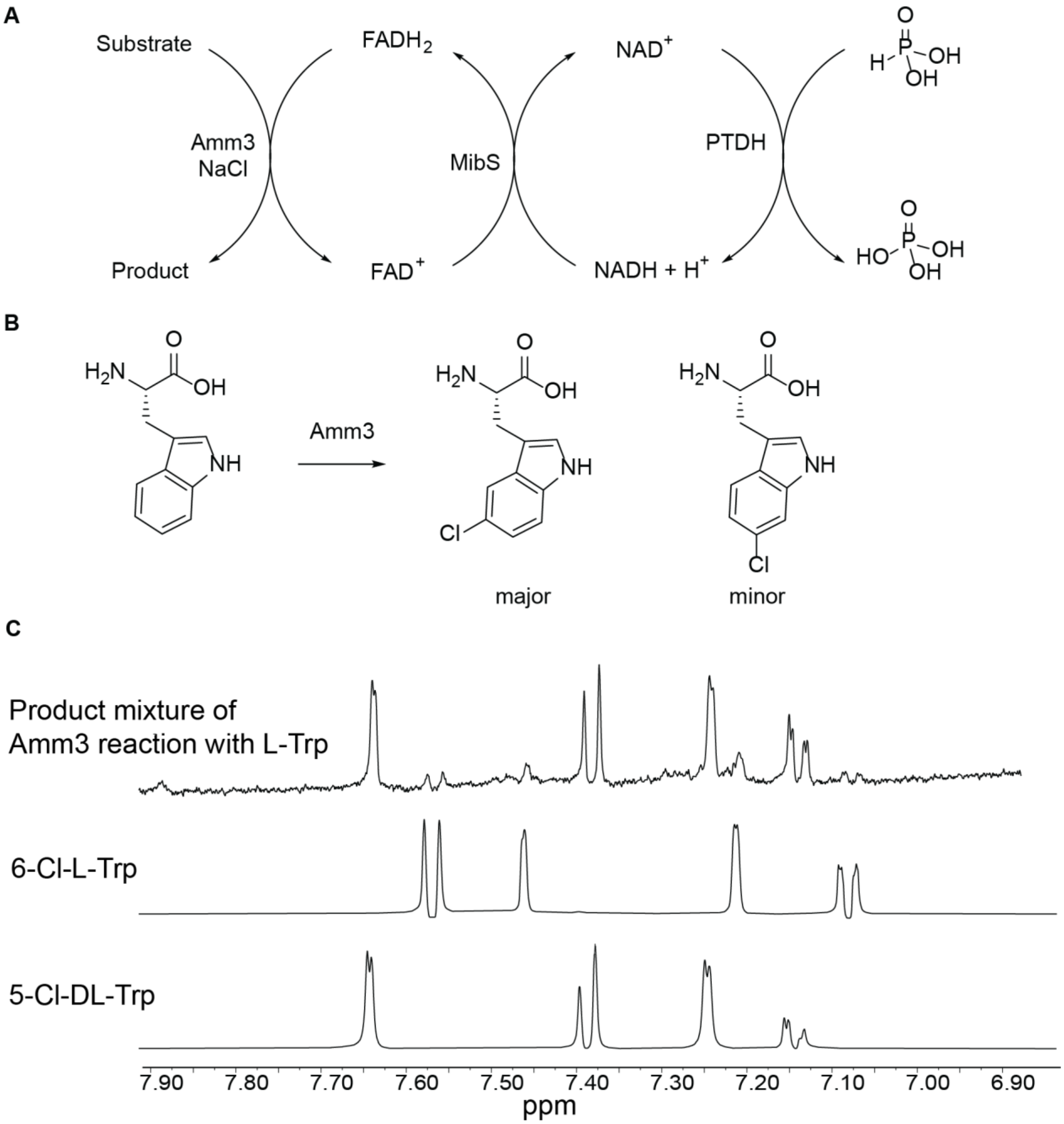


**Figure S15.** Reaction of Amm3 with Trp. (A) In vitro cascade reaction to recycle NADH as a cofactor was implemented by using phosphite dehydrogenase (PTDH) as a coupling enzyme.[4] $H_2O_2$ built up was decreased by using catalase. (B) Trp was accepted as a substrate for Amm3. Chlorination was observed mostly at C-5 of the indole, with minor C-6 chlorination. (C) Isolation of the chlorinated product of the Trp reaction with Amm3 by HPLC and analysis by $^1$H NMR revealed the major product of the reaction is 5-Cl-Trp whereas minor amounts of 6-Cl-Trp were also formed. The bottom two spectra were collected on commercial standard samples.

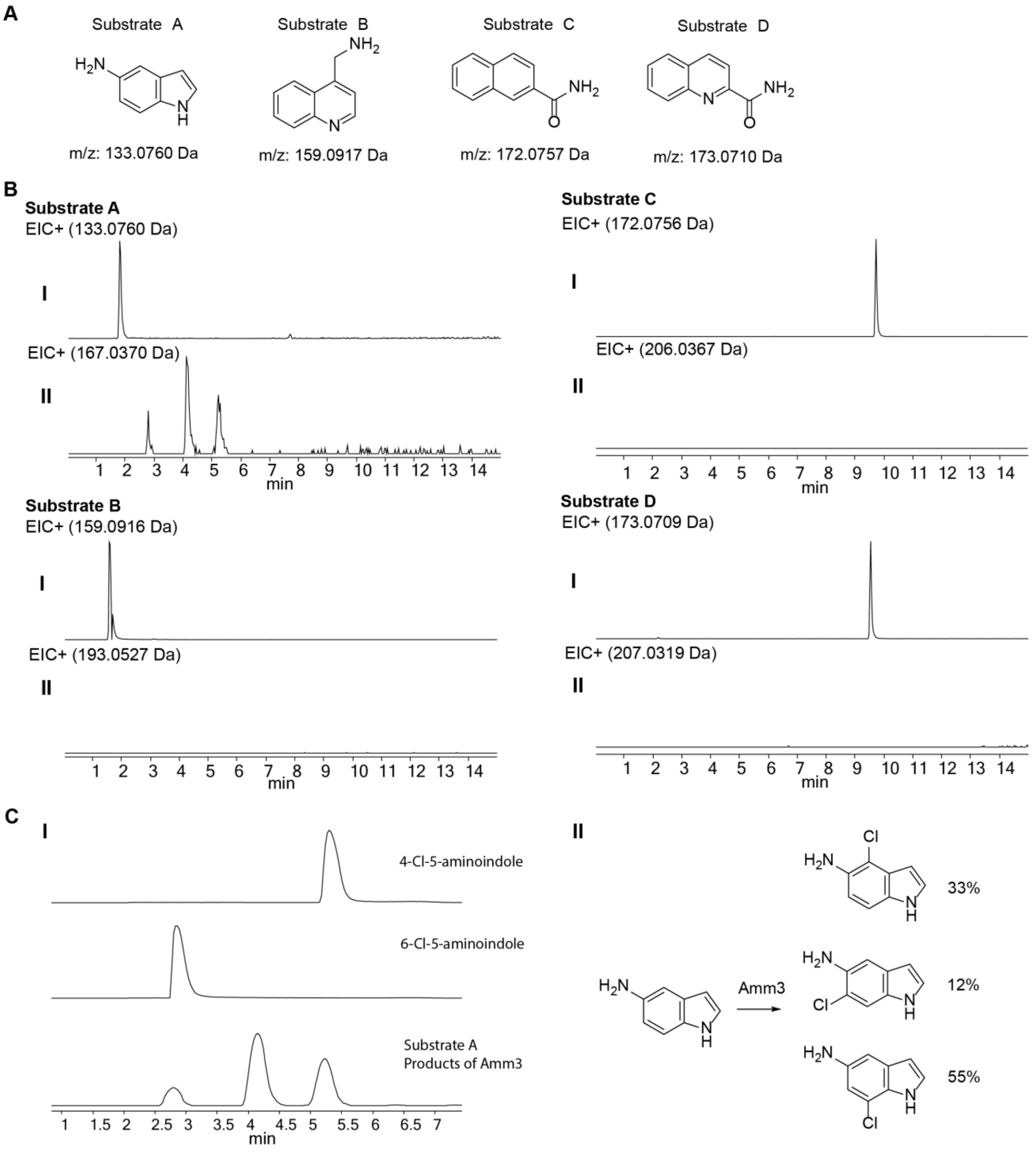


**Figure S16.** Substrate scope for chlorination catalyzed by Amm3. (A) Indole and quinoline analogs tested as potential substrates for Amm3. (B) Only the indole containing substrate A was processed by Amm3; bulkier quinoline or naphthalene containing compounds were not chlorinated by Amm3. (C) Further examination of the products of the reaction of substrate A with Amm3 by LC-MS (panel I) revealed that 4-Cl-5-aminoindole, 6-Cl-5-aminoindole, and putatively 7-Cl-5-aminoindole were produced (panel I and II).

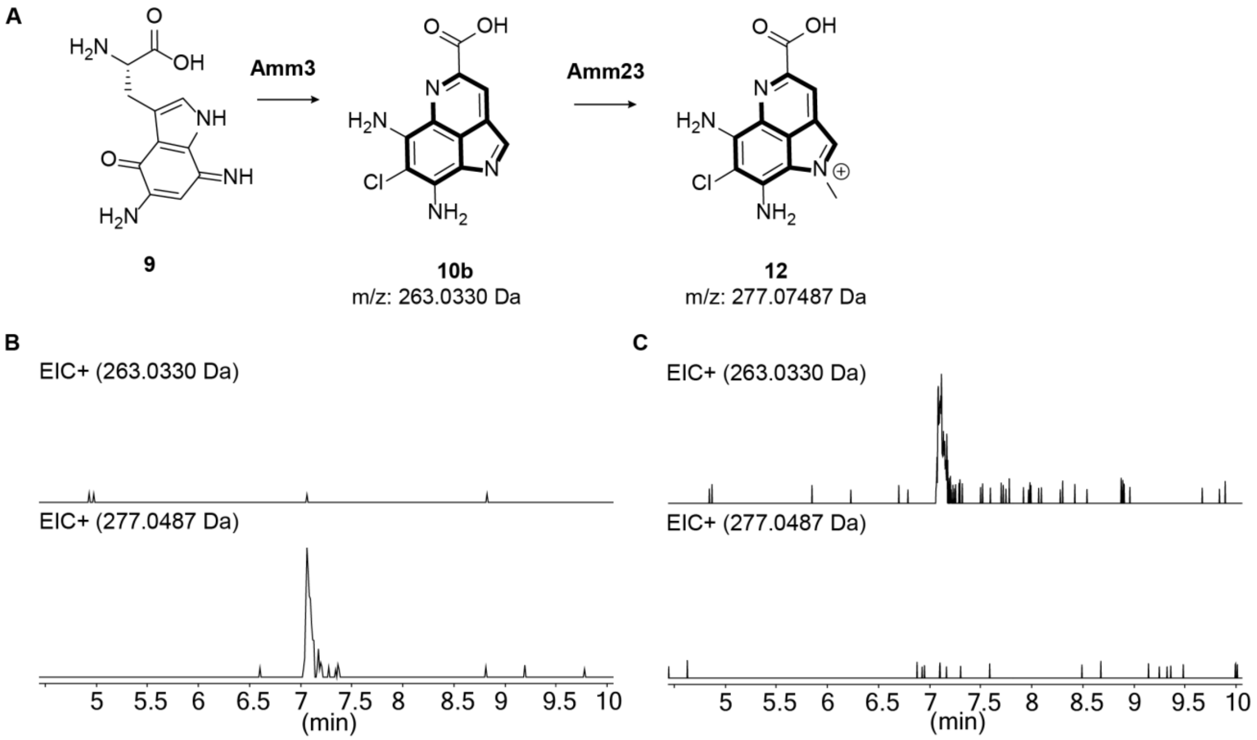


**Figure S17.** In vitro reaction of Amm23 with intermediate **10b**. (A) The product of the reaction of **9** with Amm3 produces **10b**. The reaction mixture was desalted and used in separate reactions with Amm23: (B) LC-MS analysis of a reaction containing **10b**, SAM, and Amm23. (C) LC-MS analysis of a reaction containing **10b** and SAM but without Amm23.

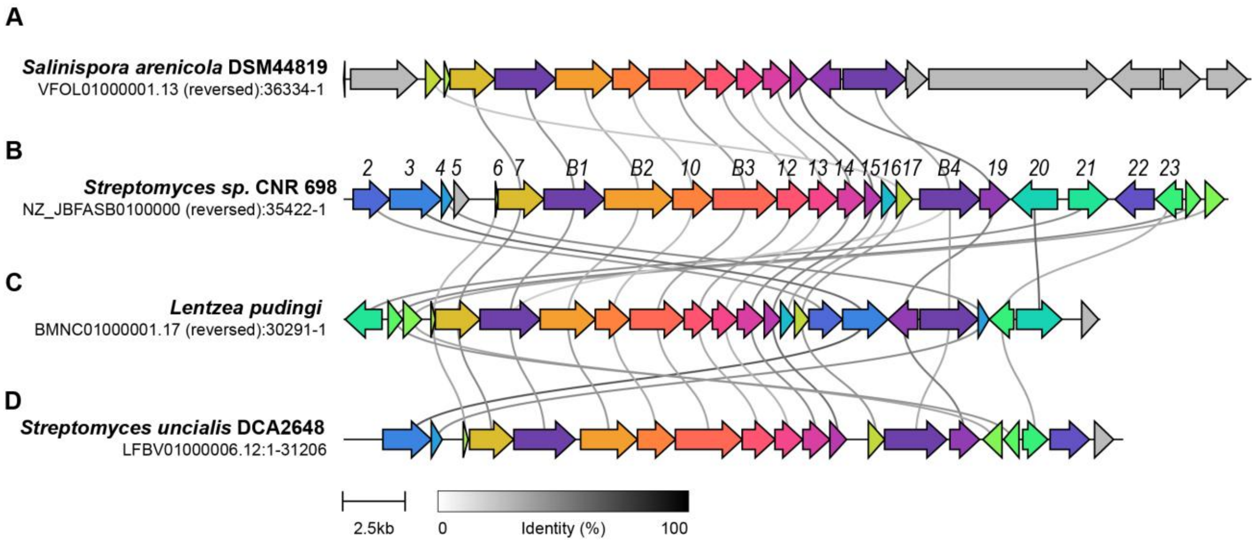


**Figure S18.** Comparison of the BGC from *Streptomyces sp.* CNR698 that produces ammosamide with the lymphostin producing BGC from *Salinospora arenicola* DSM44819, with the putative ammosamide-like-producing orthologous BGC from *Lentzea pudingi* that lacks an *amm22* gene homolog, and with the ammosester-producing BGC from *S. uncialis* DCA2648. The comparison was generated using Clinker.
